# Genomic selection strategies for clonally propagated crops

**DOI:** 10.1101/2020.06.15.152017

**Authors:** Christian R. Werner, R. Chris Gaynor, Daniel J. Sargent, Alessandra Lillo, Gregor Gorjanc, John M. Hickey

## Abstract

For genomic selection in clonal breeding programs to be effective, crossing parents should be selected based on genomic predicted cross performance unless dominance is negligible. Genomic prediction of cross performance enables a balanced exploitation of the additive and dominance value simultaneously. Here, we compared different strategies for the implementation of genomic selection in clonal plant breeding programs. We used stochastic simulations to evaluate six combinations of three breeding programs and two parent selection methods. The three breeding programs included i) a breeding program that introduced genomic selection in the first clonal testing stage, and ii) two variations of a two-part breeding program with one and three crossing cycles per year, respectively. The two parent selection methods were i) selection of parents based on genomic estimated breeding values, and ii) selection of parents based on genomic predicted cross performance. Selection of parents based on genomic predicted cross performance produced faster genetic gain than selection of parents based on genomic estimated breeding values because it substantially reduced inbreeding when the dominance degree increased. The two-part breeding programs with one and three crossing cycles per year using genomic prediction of cross performance always produced the most genetic gain unless dominance was negligible. We conclude that i) in clonal breeding programs with genomic selection, parents should be selected based on genomic predicted cross performance, and ii) a two-part breeding program with parent selection based on genomic predicted cross performance to rapidly drive population improvement has great potential to improve breeding clonally propagated crops.

## Introduction

In this paper we show that, for genomic selection in clonal breeding programs to be effective, crossing parents should be selected based on genomic predicted cross performance, unless dominance is negligible. In most plant and animal breeding programs which apply genomic selection, new parents are selected based on their genomic estimated breeding value (e.g. Meuwissen et al., 2016; Crossa et al., 2017). The genomic estimated breeding value (commonly referred to as GEBV) is by definition the sum of the average effects predicted for all marker alleles of a genotype, while dominance deviation, which cannot be directly passed on to the progeny, is not considered (Goddard, 2009; Su et al., 2012). Selection based on the genomic estimated breeding value aids breeders in increasing the frequency of alleles with beneficial additive genetic effects in a given breeding population. As a result, heterozygosity is reduced. Although selection for the genomic estimated breeding value will increase the additive value over time, it may lead to a reduction of the dominance value, unless dominance is negligible. In the long term, using the genomic estimated breeding value to select new parents in breeding programs which deliver outbred varieties, such as in clonal plant breeding programs, might not be the optimal method to use in order to maximize the total genetic value of the breeding population in a sustainable fashion.

Many major food crops, including nearly all types of fruit and all important roots and tubers, are clonally propagated (Grüneberg et al., 2009; Bradshaw, 2016). In clonal breeding programs, new genotypes are created by sexual reproduction and multiplied through clonal propagation (Bisognin, 2011; Gemenet and Khan, 2017). The new genotypes are first tested as seedlings in unreplicated trials during the initial phase of the breeding program. Clonal propagation creates genetically identical plants from selected seedlings, which enables the testing of genotypes in clonal plots, using multiple replications, environments and years.

Breeders use multiple stages of testing to identify and select the best genotypes in their breeding population. As the testing progresses, the number of genotypes is successively reduced and those remaining are tested more intensively at increasingly higher numbers. The selected genotypes are used to achieve two specific objectives:

i. Generation of an improved offspring population via recombination of selected parents.
ii. Release of the best performing genotypes as improved clonal varieties.

The time from recombination to the release of an improved clonal variety spans several years. Traditionally, selection is based on phenotypic performance and the next generation’s parents are selected in the later testing stages of the breeding program, which results in a long generation interval (Bradshaw, 2016), even in species with short generational times, such as strawberry.

Genomic selection offers great potential to optimize the process of identification of the best clones for varietal development, as well as the selection of new crossing parents. Genomic selection exploits associations between genomic markers and phenotypes to predict the value of genotypes based on their genomic marker profiles (Goddard and Hayes, 2007). The implementation of genomic selection provides three key advantages:

i. The generation interval can be reduced, since new parents can be selected as soon as they are genotyped.
ii. The selection accuracy can be increased, especially in early testing stages of a breeding program where the number of replications and environments is low.
iii. The selection intensity can be increased, for example by genotyping and predicting more genotypes than could be tested in the field.

These advantages allow for several opportunities to reorganize conventional breeding programs. For example, in the context of breeding programs to develop inbred lines, Gaynor et al. (2017) presented a two-part breeding program employing genomic selection, which reorganized a plant breeding program into:

i. A population improvement component to develop improved germplasm through rapid recurrent genomic selection, and
ii. A product development component to identify the best performing genotypes for varietal development.

In stochastic simulation, the two-part breeding program doubled the rate of genetic gain relative to a conventional breeding program without increasing cost.

In a clonal breeding program, the reorganization in two parts combined with genomic selection would allow breeders to minimize the generation interval and could substantially increase selection accuracy at the seedling stage.

The generation interval could be reduced to a year or even less since new parents can be selected as soon as the seedlings are genotyped. For example, the generation interval in conventional strawberry breeding programs can be four to five years due to the time it takes for testing to generate sufficient phenotypic records to accurately assess a genotype. Genomic selection applied in the seedling stage could result in up to five times the genetic gain achieved in a conventional strawberry breeding program in the same amount of time if the impact of the three other factors in the breeder’s equation (i.e., selection intensity, diversity and selection accuracy) remained constant.

The selection accuracy in the seedling stage could be increased since genomic selection allows seedlings to be selected based on their predicted performance as clones instead of their phenotypic performance *per se*. This is achieved when the genomic selection model to select seedlings is trained using clonal phenotypes. In clonal breeding programs, the seedling stage represents a severe genetic bottleneck; in conventional strawberry breeding programs only a few hundred genotypes among 10,000 – 20,000 unreplicated seedlings are selected and tested as clones. Selection accuracy is extremely low at the seedling test stage for three reasons (Grüneberg et al., 2009), which are:

i. Seedlings and clones with the same genotype can differ in their morphology and performance.
ii. Seedlings and clones are often grown in different environments. For example, in European strawberry breeding programs, seedlings are grown in matted rows on the soil and clones are grown as single pot plants on highly controlled table top systems.
iii. Single plant assessment of mostly general appearance and/or a few key traits in the seedling stage shows low heritability and has low correlation with the breeding goal trait (e.g., yield).

Replacing phenotypic selection in the seedling stage with genomic selection based on the predicted performance as clones eliminates all three challenges in one step. It also allows for early evaluation of important traits that are typically not evaluated until later testing stages of the breeding program, e.g. flavour and shelf life.

In clonally propagated crops, however, dominance may affect the performance of breeding programs which implement genomic selection. The genotypes in clonally propagated crops are typically heterozygous. The genetic value of heterozygous genotypes is a function of additive and non-additive gene action (Falconer and Mackay, 1996). If, for the sake of simplicity, epistasis is ignored, the non-additive gene action is entirely defined by dominance. Whilst the differences in the genetic values between genotypes are based on both additive and non-additive genetic effects, the additive genetic variation is the crucial component which defines long-term genetic gain in a breeding population subjected to recurrent selection (Bradshaw 2016). Hence, breeders face the challenging task of having to increase the additive value over time while simultaneously maintaining the dominance value via selection and recombination of the best parents. The relative importance of these two targets is a function of the dominance degree at the loci affecting the trait under consideration, which is mostly unknown.

We hypothesise that genomic prediction of cross performance is a better method to select new parents in a clonal breeding program than using the genomic estimated breeding value. When genomic prediction of cross performance is used, pairs of parents are selected based on the expectation of the total genetic value of their progeny. Genomic prediction of cross performance could allow breeders to simultaneously increase the frequency of alleles with beneficial additive effects and maintain heterozygosity in the population to exploit dominance effects. In the long term, using genomic prediction of cross performance to select new parents in a clonal breeding program could be an effective method to sustainably maximize the total genetic value of the breeding population.

To test our hypothesis, we used stochastic simulation to evaluate three breeding programs and two parent selection methods to deploy genomic selection in breeding clonally propagated crops under different dominance degrees. The three breeding programs included:

i. A breeding program that introduced genomic selection in the first clonal testing stage, and
ii. Two variations of a two-part breeding program (Gaynor et al., 2017) with one and three crossing cycles per year, respectively.

The two parent parental selection methods were:

i. Selection of parents based on genomic estimated breeding values, and
ii. Selection of parents based on genomic predicted cross performance.

The six combinations of breeding program and parent selection method were compared to a conventional breeding program using phenotypic selection.

We observed that the breeding programs using selection of parents based on genomic predicted cross performance produced faster genetic gain than parent selection based on genomic estimated breeding values unless dominance was negligible. The highest rates of genetic gain were generated by the two-part breeding programs with parent selection based on genomic predicted cross performance.

## Materials and methods

Stochastic simulations were used to evaluate six combinations of three breeding programs and two parent selection methods to deploy genomic selection in breeding clonally propagated crops with diploid (-like) meiotic behaviour. Therefore, we simulated a quantitative trait representing yield under four different dominance degrees and evaluated the long-term efficacy of the six combinations of breeding programs and parent selection methods compared to a conventional breeding program using phenotypic selection.

The material and methods are subdivided into two sections. The first section describes the simulation of the founder genotype population and the second section describes the simulation of the breeding programs.

The simulation of the founder genotype population comprised:

i. Genome simulation: a heterozygous genome sequence was simulated for a hypothetical diploid and clonally propagated crop species.
ii. Simulation of founder genotypes: the simulated genome sequences were used to generate a base population of 60 diploid founder genotypes.
iii. Simulation of genetic values: A single trait representing yield was simulated for all founder genotypes by summing the additive and dominance effects at 20,000 quantitative trait nucleotides. Four different dominance degrees were simulated including 0, 0.1, 0.3 and 0.9
iv. Simulation of phenotypes: Phenotypes for yield were simulated for all founder genotypes by adding random error to the total genetic value of a genotype.

The simulation of the breeding programs comprised:

i. Recent (burn-in) breeding phase: a conventional phenotypic selection breeding program for clonally propagated crops was simulated for a period of 20 years (burn-in) to provide a common starting point for the future breeding phase.
ii. Future breeding phase: six combinations of three breeding programs and two parent selection methods to deploy genomic selection in clonally propagated crops were simulated and compared to the conventional breeding program for 20 years of breeding. In detail, we describe:

a. The genomic selection model used for genomic prediction.
b. The two parent selection methods including parent selection based on genomic estimated breeding values and parent selection based on genomic predicted cross performance.
c. The three breeding programs with genomic selection including a breeding program which implemented genomic selection in the clonal testing stage 1, and two variations of a two-part breeding program which implemented genomic selection in the seedling stage with one and three crossing cycles per year, respectively.
d. Comparison of the breeding programs based on the mean total genetic value in clonal testing stage 1.

### Simulation of the founder genotype population

#### Genome simulation

A heterozygous genome sequence was simulated for each genotype of a hypothetical diploid and clonally propagated crop species. The simulated genome consisted of 20 chromosome pairs with a physical length of 10_8_ base pairs and a genetic length of 100 centiMorgans (cM), resulting in a total genetic length of 2,000 cM comparable to that of the *Fragaria × ananassa* genome (Sargent et al., 2009, 2016; van Dijk et al., 2014; Bassil et al., 2015). The chromosome sequences were generated using the Markovian coalescent simulator (MaCS; Chen et al. 2009), which was deployed using AlphaSimR version 0.11.0 (Gaynor et al., 2019). Recombination rate was derived as ratio between genetic length and physical genome length (i.e., 100 cM / 10_8_ base pairs = 10_−8_). The per-site mutation rate was set to 2.5 × 10_−8_ mutations per base pair. Effective population size (N_e_) was set to 100 and resulted from a simulated coalescence process with an effective population size of 500, 1,250, 1,500, 3,500, 6,000, 12,000 and 100,000 set for 100, 500, 1,000, 5,000, 10,000, and 100,000 generations ago. Successive reduction of the effective population size was used to reflect a progressive restriction of genetic variation due natural and artificial selection.

#### Simulation of founder genotypes

The simulated genome sequences were used to generate a base population of 60 diploid founder genotypes in Hardy-Weinberg equilibrium. These genotypes were formed by randomly sampling 20 chromosome pairs per genotype and served as initial parents in the burn-in phase. A set of 1,000 biallelic quantitative trait nucleotides (QTN) and 1,000 single nucleotide polymorphisms (SNP) were randomly sampled along each chromosome to simulate a quantitative trait that was controlled by 20,000 QTN and a SNP marker array with 20,000 markers.

#### Simulation of genetic values

Genetic values for a single trait representing yield were simulated by summing the genetic effects at the 20,000 randomly sampled QTN. Three types of biological effects were modelled at each QTN to simulate genetic values: additive effects, dominance effects and genotype-by-environment effects. Under the AlphaSimR framework, this is referred to as an ADG trait. We will give only a brief summary of the modelling procedure, while a detailed description can be found in the vignette of the AlphaSimR package (Gaynor et al., 2019).

Additive effects (*a*) were sampled from a standard normal distribution and scaled to obtain an additive variance of 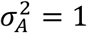 in the founder population. Genotype-by-environment effects were modelled using an environmental covariate and a genotype-specific slope. The environmental covariate represented the environmental component of the genotype-by-environment interaction and was sampled for each year of the simulation from a standard normal distribution. The genotype-specific slope represented the genetic component of the genotype-by-environment interaction. The effects for the genotype specific slope were sampled from a standard normal distribution and scaled to obtain a genotype-by-environment interaction variance of 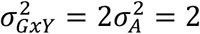 in the founder population.

Dominance effects (*d*) for all QTN were calculated by multiplying the absolute value of its additive effect *a*_*i*_ by a locus-specific dominance degree *δ*_*i*_. A dominance degree of 0 represents no dominance and a dominance degree of 1 represents complete dominance. Dominance degrees between 0 and 1 correspond to partial dominance, and values above 1 correspond to over-dominance. Dominance degrees were sampled from a normal distribution with mean dominance coefficient *μ*_*δ*_ and variance 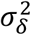:

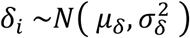

The dominance effect of QTN *i* was calculated as:

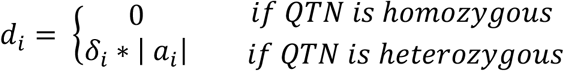

Three levels of average dominance degrees, 0.1, 0.3 and 0.9, were used to simulate positive directional dominance and compared to zero dominance (i.e., additive genetic control). The variance 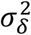 was set to 0.2. The dominance variance 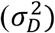 was then calculated based on the simulated dominance effects.

#### Simulation of phenotypes

Phenotypes for yield were generated by adding random error to the genetic value of a genotype. The random error was sampled from a normal distribution with mean zero and an error variance 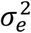 defined by the target level of heritability at each testing stage of the breeding program. In the founder population, entry-mean values for narrow-sense heritability (*h*^2^) were set to 0.1 in the seedling stage and to 0.3 in clonal testing stage 1 of the breeding program, with 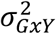 set to 0. Entry-mean levels for narrow-sense heritabilities in later testing stages increased as a result of an increased number of replicates per genotype and are shown in Table 1. Narrow-sense heritabilities were calculated using the following equation:

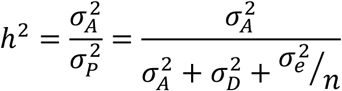

**Table 1.** Number of tested genotypes, replications and heritabilities used in the conventional breeding program

| Year | Stage | Tested genotypes | Reps | Narrow-sense heritability ($h_2$)* |
| --- | --- | --- | --- | --- |
| 1 | Seedlings | 15,000 | 1 | 0.10 |
| 2 | Clonal stage 1 | 1,000 | 1 | 0.30 |
| 3 | Clonal stage 2 | 100 | 2 | 0.46 |
| 4 | Clonal stage 3 | 20 | 4 | 0.63 |
| 5 | Clonal stage 4 | 5 | 6 | 0.72 |
| 6 | Clonal stage 5 | 5 | 6 | 0.72 |

\*entry-mean values based on the $\sigma_A^2 : \sigma_P^2$ ratio in the founder population

### Simulation of the breeding programs

#### Recent (burn-in) breeding phase

A conventional breeding program for clonally propagated crops employing phenotypic selection was simulated for a period of 20 years (burn-in) to provide a common starting point for the future breeding phase. Each year of the conventional breeding program started with a crossing block of 60 parental genotypes. These genotypes were crossed to generate new seedlings, followed by a six year evaluation period that involved six stages of testing. Selection of new parents and selection of the best clones in each testing stage were based on phenotypic records. The structure and the values for key parameters of the conventional breeding program were guided by a commercial strawberry breeding program in the United Kingdom. Table 1 presents the number of tested genotypes and replications for each testing stage of the conventional breeding program as shown in Figure. 1.

**Figure 1.**
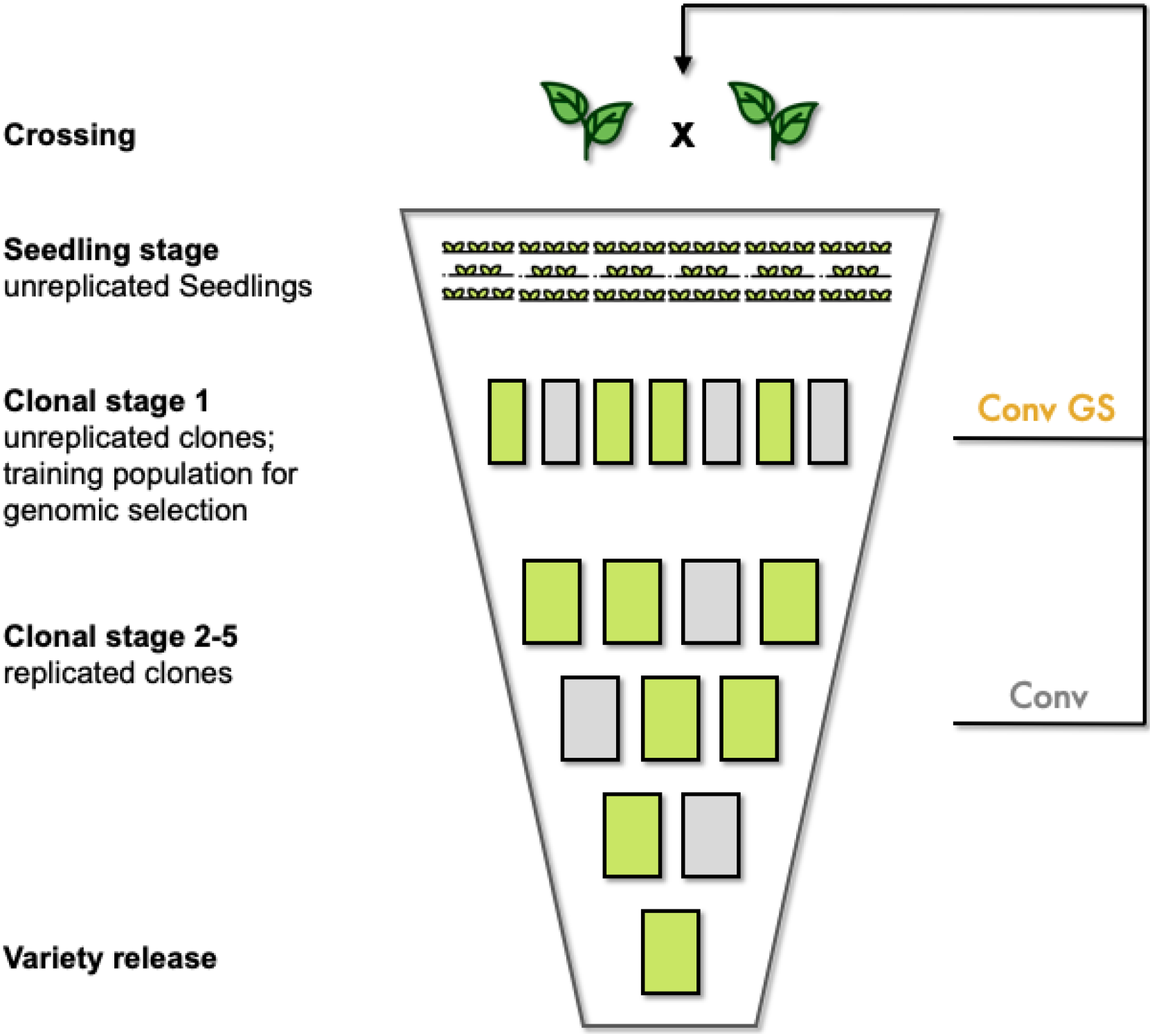
Schematic overview of the conventional breeding program and the conventional breeding program with genomic selection. The conventional breeding program (Conv) was used in the burn-in breeding phase and served as a control in the future breeding phase. In the conventional breeding program, parents were selected in clonal stages 2-5. The conventional breeding program with genomic selection reduced the generation interval to two years by selecting parents in clonal stage 1 based on either genomic estimated breeding values or genomic predicted cross performance. The genotypes in clonal stage 1 served as training population.

In order to fill the breeding pipeline and generate a starting point for the burn-in phase, six cycles of crossing and selection were conducted prior to the burn-in phase. Each of these six cycles started with the same 60 founder genotypes to generate 150 F_1_-families with 100 seedlings each, using random sampling of bi-parental crosses without replacement. Starting from the set of 15,000 seedlings after the first crossing cycle, the best genotypes were advanced one stage per cycle using phenotypic selection until each testing stage was filled with a set of genotypes. Replacement of parents was omitted during the filling of the breeding pipeline. This was done to ensure that total genetic variance in the founder genotypes remained unchanged until the actual burn-in phase started.

In the burn-in phase, selection of new parents was carried out in the clonal testing stages 2, 3, 4 and 5. Each year, the 30 genotypes in the crossing block with the poorest *per se* performance were replaced by new parents. At first, all 30 genotypes in the clonal testing stages 3, 4 and 5 were added to the crossing block as new parents if they were not already represented. The remaining free slots in the crossing block were filled with the best genotypes from the clonal testing stage 2.

#### Future Breeding Phase

The future breeding phase was used to evaluate six combinations of two breeding programs and two parent selection methods to deploy genomic selection in clonally propagated crops under different dominance degrees. These six combinations were simulated for an additional 20 years of breeding and compared to the conventional breeding program. The two genomic selection breeding programs included a conventional breeding program with genomic selection which introduced genomic selection in clonal testing stage 1 (Fig. 1), and two variations of a two-part breeding program which introduced genomic selection in the seedling stage with one and three crossing cycles per year, respectively (Fig. 2). The two parent selection methods were selection of new parents based on genomic estimated breeding values, and selection of new parents based on genomic predicted of cross performance. In order to obtain approximately equal annual operating costs, the number of seedlings was reduced in the two breeding programs with genomic selection to compensate for the additional costs of genotyping. Estimated costs were set to $20 for phenotypic evaluation and $25 for array genotyping per genotype after consultation with strawberry breeders. Table 2 shows the number of crosses and seedlings per year for the conventional breeding program and the three breeding programs with genomic selection.

**Table 2.**
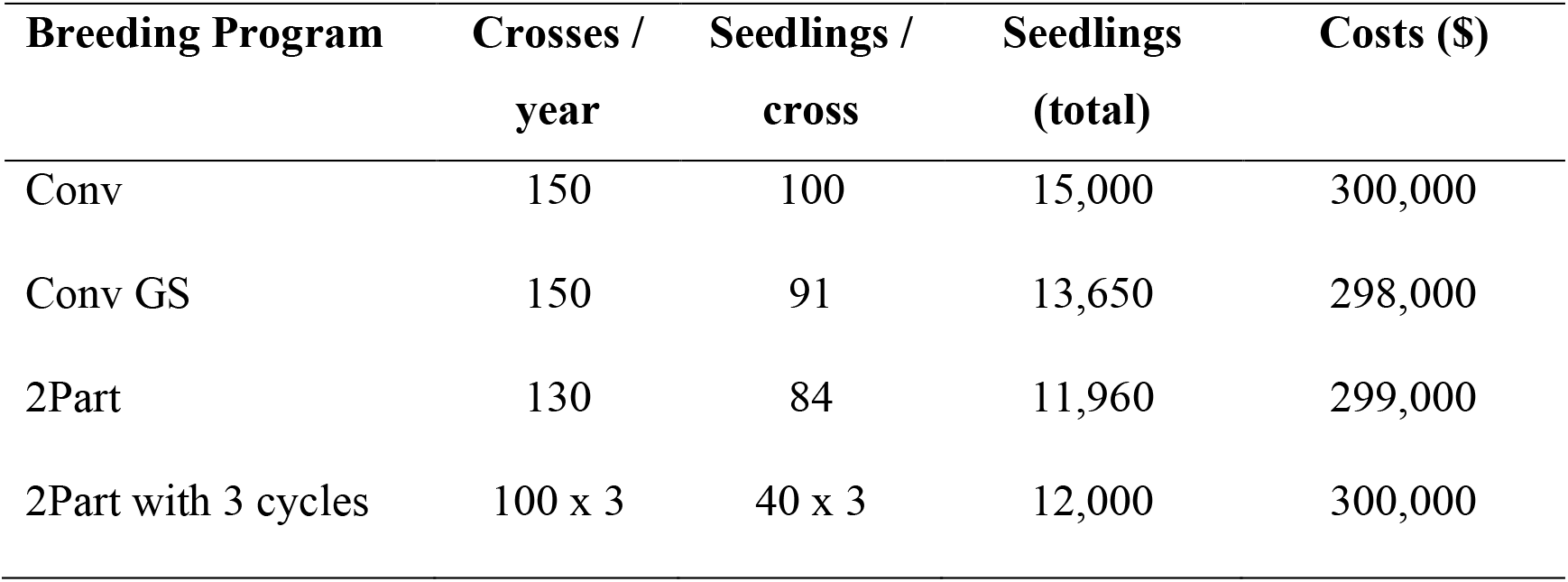
Number of crosses per year and seedlings per cross, total number of seedlings and annual costs of the simulated breeding programs (Conv, conventional breeding program; Conv GS, conventional breeding program with genomic selection; 2Part, two-part breeding program)

**Figure 2.**
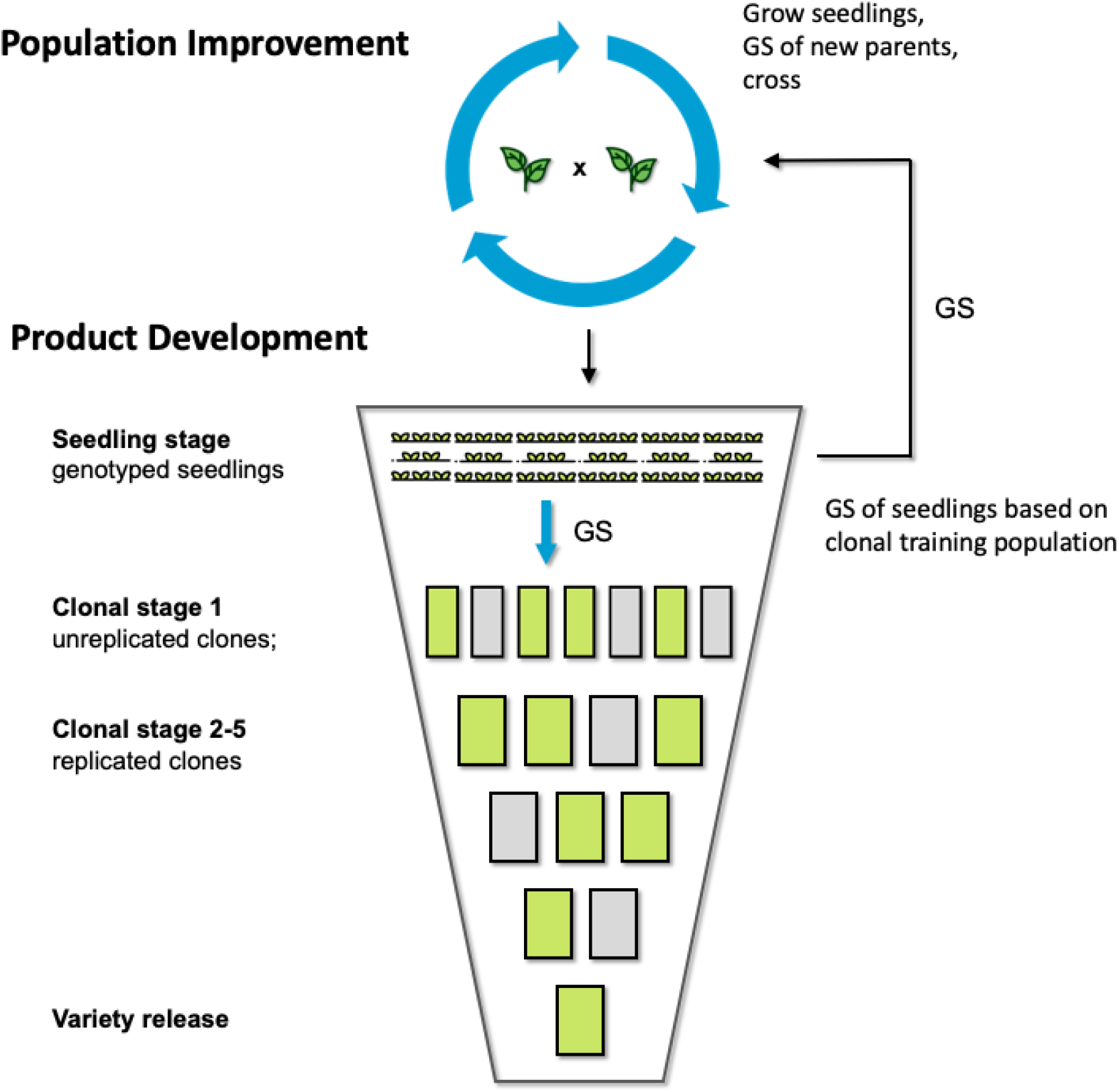
Schematic overview of the two-part breeding program. The two-part breeding program reorganized the conventional breeding program into i) a population improvement component to develop improved germplasm through rapid recurrent genomic selection; and ii) a product development component to identify the best performing genotypes. The population improvement component allows to have multiple cycles of crossing and selection per year before the seedlings are advanced to the product development component based on their genomic estimated genetic values. New parents during population improvement were selected based on either genomic estimated breeding values or genomic predicted cross performance.

#### Genomic Selection Model

Genomic predictions were calculated using a ridge regression model (RR-BLUP) including year as a fixed effect, additive and dominance SNP effects, and a covariate accounting for directional dominance (or inbreeding depression) based on average individual heterozygosity as described in detail by Xiang et al. (2016). The effect estimated for the covariate accounting for directional dominance was divided by the number of SNPs and added to the SNP-specific dominance effects. To obtain genomic estimated breeding values, the predicted additive and dominance SNP effects at each marker locus were used to calculate the average effect of an allele substitution for each SNP (Varona et al., 2018), and all the substitution effects were summed. To obtain genomic estimated genetic values, the predicted additive and dominance SNP effects at each marker locus were summed. The initial training population at the start of the future breeding phase consisted of all the genotypes from clonal testing stage 1 of the last three years of the burn-in phase. The training population included 3,000 genotypes and 3,220 phenotypic records. In every year of the future breeding phase, 1,000 new genotypes from clonal testing stage 1 were added to the training population.

#### Parent selection methods

Two parent selection methods were compared for the selection and crossing of new parents in the two breeding programs with genomic selection. The first parent selection method will be referred to as *parent selection based on genomic estimated breeding values*. This method represented a conventional “good by good” crossing scheme. The genotypes with the highest genomic estimated breeding values were selected as new parents and used to completely replace the previous year’s crossing block. Crossing was implemented as random sampling of bi-parental combinations without replacement. The second parent selection method will be referred to as *parent selection based on genomic predicted cross performance*. This method implemented systematic selection of bi-parental crosses. The best bi-parental crosses were selected based on the predicted mean genetic values of the F_1_ of a cross. In this way, the average amount of heterosis predicted for the F_1_ due to complementarity between two parents was directly considered in the parent selection process. The mean genetic value of the F_1_ of a cross was predicted using the following equation given by Falconer & Mackay (1996):

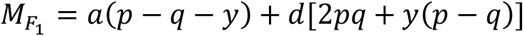

with 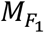 being the predicted mean genotypic value of the F_1_, *a* and *d* being the additive and dominance effects of the SNP markers, *p* and *q* being the marker allele frequencies of one parent and *y* representing the difference of gene frequency between the two parents. The concept of the crossing block was abandoned.

#### Conventional breeding program with genomic selection

The conventional breeding program with genomic selection introduced genomic selection in clonal testing stage 1. The structure of the conventional breeding program with genomic selection is shown in Figure 1. All 1,000 genotypes in clonal testing stage 1 were genotyped and phenotyped to serve as the training population for the genomic selection model. When parents were selected based on genomic estimated breeding values, each year the best 60 genotypes in clonal testing stage 1 were used to replace the whole crossing block. When parents were selected based on genomic predicted cross performance, bi-parental cross performance was predicted for all pairwise combinations between the genotypes in clonal testing stage 1. The generation interval was two years. Genomic selection was also used to advance the best 100 clones from clonal testing stage 1 to clonal testing stage 2 based on their genomic estimated genetic value.

#### Two-part breeding programs

The two-part breeding programs reorganized the conventional breeding program into a population improvement component to develop improved germplasm through rapid recurrent genomic selection, and a product development component to identify the best performing genotypes. Two variations of the two-part breeding program with one and three crossing cycles per year respectively were simulated. The structure of the two-part breeding programs is shown in Figure 2. Genomic selection was introduced in the seedling stage. All seedlings were genotyped and phenotypic selection in the seedling stage was entirely replaced by genomic selection. All 1,000 genotypes in clonal testing stage 1 were genotyped and phenotyped to serve as the training population for the genomic selection model. Thus, a key feature of the two-part breeding program is that seedlings were selected using a prediction model that was trained with phenotypic records from clones. When parents were selected based on genomic estimated breeding values, in each crossing cycle the best 60 seedlings were used to replace the whole crossing block. When parents were selected based on genomic predicted cross performance, bi-parental cross performance was predicted for all pairwise combinations between the seedlings. The generation interval was one year with one crossing cycle per year and 1/3 year with 3 crossing cycles per year. Genomic selection was also used to advance the best 1,000 seedlings to clonal testing stage 1 and the best 100 clones from clonal testing stage 1 to clonal testing stage 2 based on their genomic estimated genetic value.

#### Comparison of the breeding programs

The performance of the three breeding programs and the two parent selection methods in comparison to the conventional breeding program was evaluated by measuring the mean total genetic value in clonal testing stage 1. Each evaluation included ten simulation runs. The mean total genetic value was measured in clonal testing stage 1 for two reasons:

i. It was the earliest testing stage in which clones were evaluated.
ii. The general trends observed for genetic gain in clonal testing stage 1 were representative for genetic gain in the seedling stage and genetic gain in later testing stages of the breeding programs.

The additive value, the dominance value and the genomic inbreeding coefficient over time were also measured in clonal testing stage 1. The genomic inbreeding coefficient was calculated as the percentage increase of homozygosity at all quantitative trait nucleotides relative to the average homozygosity observed in the founder population.

All breeding programs were compared for total genetic variance, additive variance and dominance variance over time, results are shown in the supplementary material (Fig. S1-S3).

Prediction accuracy was assessed as the Pearson correlation coefficient in two different ways:

i. Prediction accuracy was assessed in the three breeding programs with genomic selection as the accuracy of the parent selection method including parent selection based on genomic estimated breeding values and parent selection based on genomic predicted cross performance.
ii. Prediction accuracy was assessed as the prediction accuracy of the total genetic value in the seedling stage, which was used to advance seedlings to clonal testing stage 1.

## Results

The results show that for genomic selection in a clonal breeding program to be effective, crossing parents should be selected based on genomic predicted cross performance unless dominance is negligible. Selection of parents based on genomic predicted cross performance produced faster genetic gain than selection of parents based on genomic estimated breeding values when the dominance degree was greater than zero (Fig. 3). As the dominance degree increased, selection of parents using genomic prediction of cross performance also produced increasingly more genetic gain than selection based on genomic estimated breeding values. The two variations of the two-part breeding program using genomic prediction of cross performance always produced the most genetic gain unless dominance was negligible. However, while the two-part breeding program with three crossing cycles per year produced the most genetic gain when the dominance degree was low, the two-part breeding program with one crossing cycle per year produced the most genetic gain when the dominance degree was high. The breeding programs using selection of parents based on genomic estimated breeding values on the other hand, produced negative genetic gain when the dominance degree was high. Selection of parents based on genomic prediction of cross performance was advantageous over selection of parents based on genomic estimated breeding values because it substantially reduced inbreeding in the breeding population when the dominance degree increased (Fig. 4). This enabled a better exploitation of the additive value and the dominance value simultaneously, which became more important as the dominance degree increased (Fig. 5). Additionally, selection of parents based on genomic prediction of cross performance became more accurate and selection of parents based on genomic estimated breeding values became less accurate at higher dominance degrees (Fig. 6).

**Figure 3.**
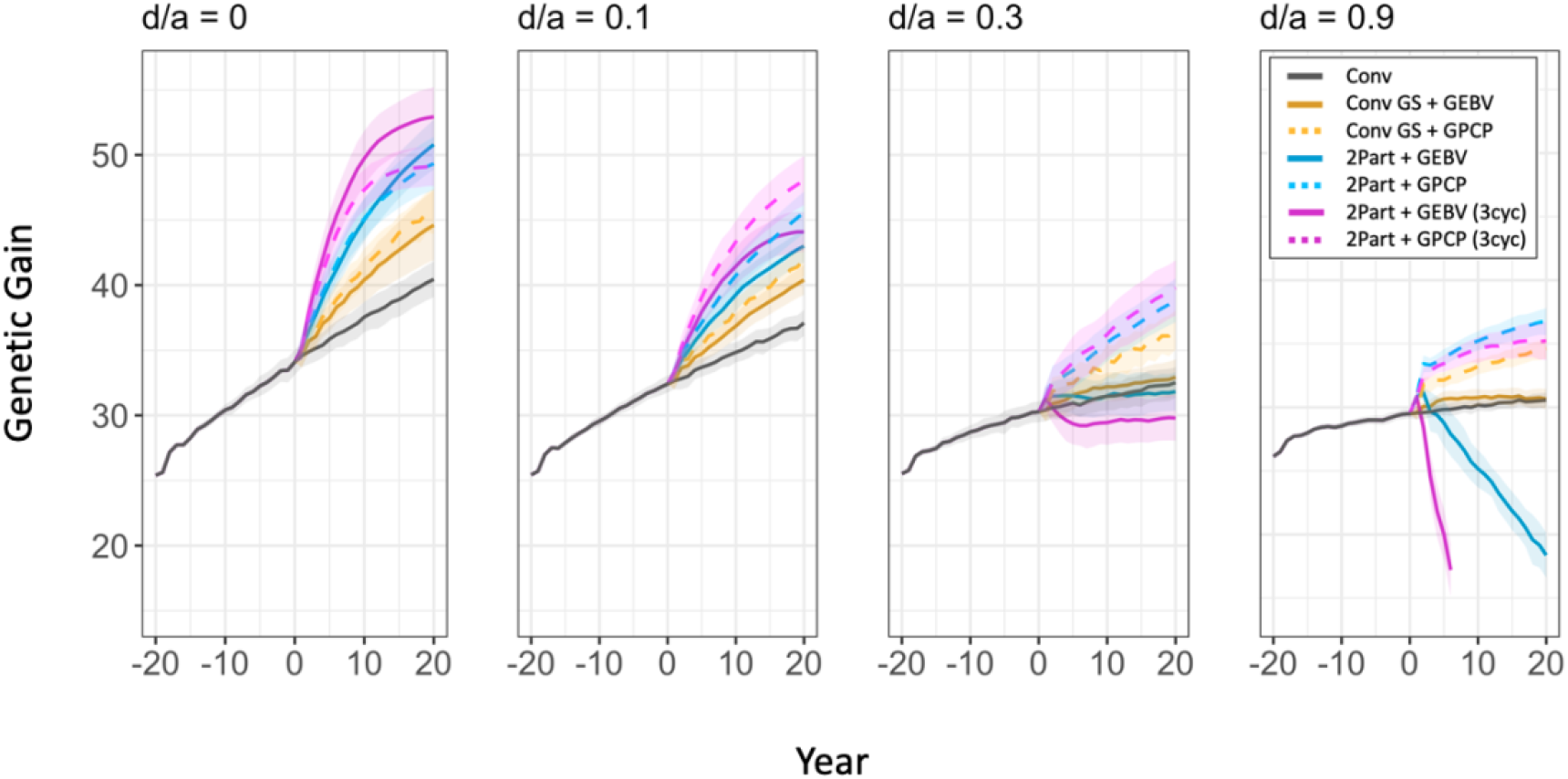
Genetic gain of the simulated breeding programs under different dominance degrees (d/a). In each panel, genetic gain is plotted as mean genetic value in clonal stage 1 for the entire burn-in breeding phase and the future breeding phase. Each line shows the mean genetic value for the 10 simulated replications and the shading shows the 95% confidence intervals. The different types of breeding program are shown in different colours. The conventional breeding program (Conv) is gray. The conventional breeding program with genomic selection (Conv GS) is yellow. The two-part breeding program with genomic selection (2Part) is shown in blue with one crossing cycle per year and in purple with three crossing cycles per year. The two types of parent selection were shown in different line-styles. Selection based on Genomic Estimated Breeding Value (GEBV) is shown by continuous lines. Selection based on Genomic Prediction of Cross Performance (GPCP) is shown by dashed lines.

**Figure 4.**
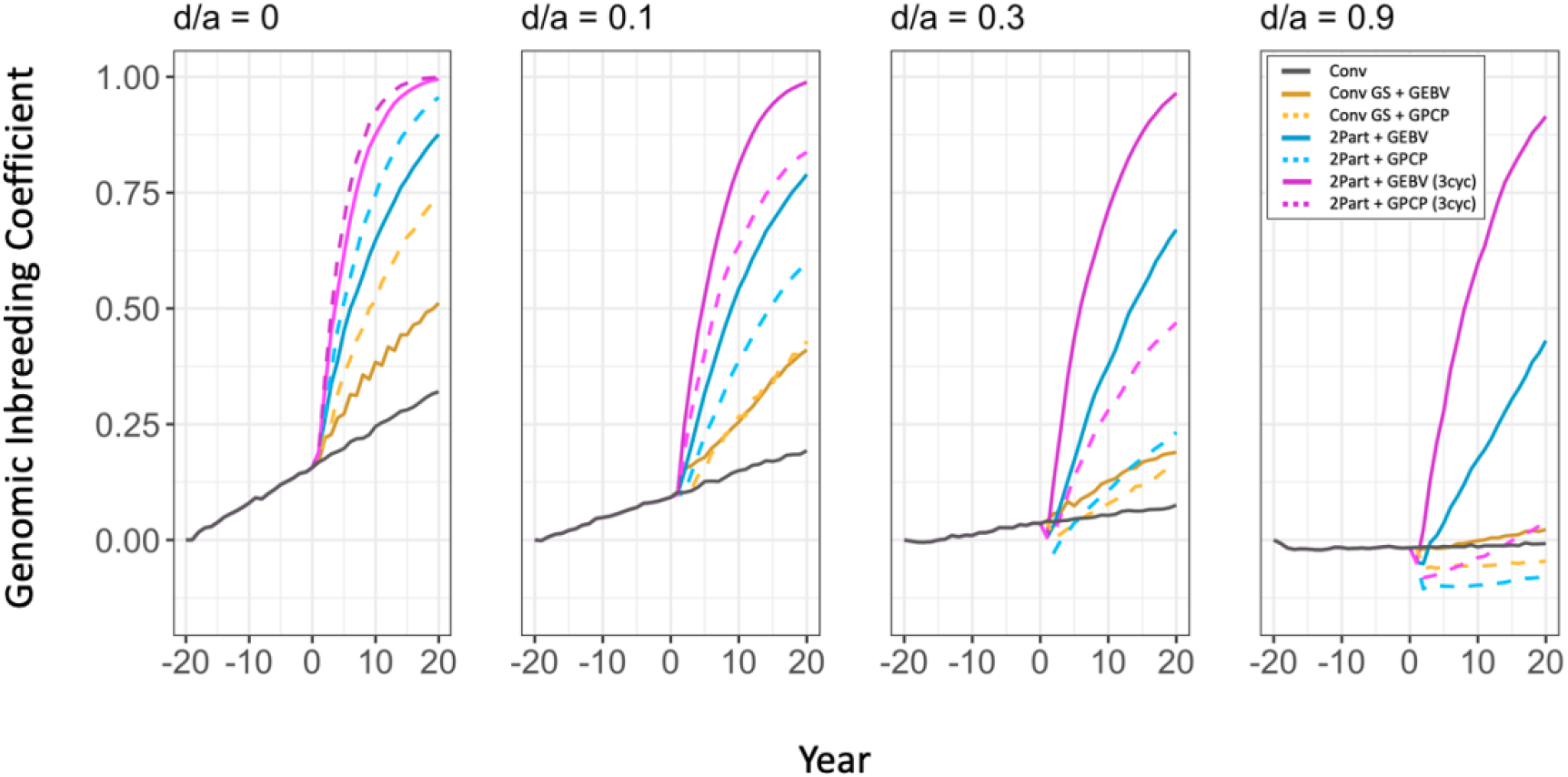
Genomic inbreeding coefficient of the simulated breeding programs under different dominance degrees (d/a). In each panel, the genomic inbreeding coefficient is plotted in clonal stage 1 for the entire burn-in breeding phase and the future breeding phase. Each line shows the mean genomic inbreeding coefficient for the 10 simulated replications. The different types of breeding program are shown in different colours. The conventional breeding program (Conv) is gray. The conventional breeding program with genomic selection (Conv GS) is yellow. The two-part breeding program with genomic selection (2Part) is shown in blue with one crossing cycle per year and in purple with three crossing cycles per year. The two types of parent selection were shown in different line-styles. Selection based on Genomic Estimated Breeding Value (GEBV) is shown by continuous lines. Selection based on Genomic Prediction of Cross Performance (GPCP) is shown by dashed lines.

**Figure 5.**
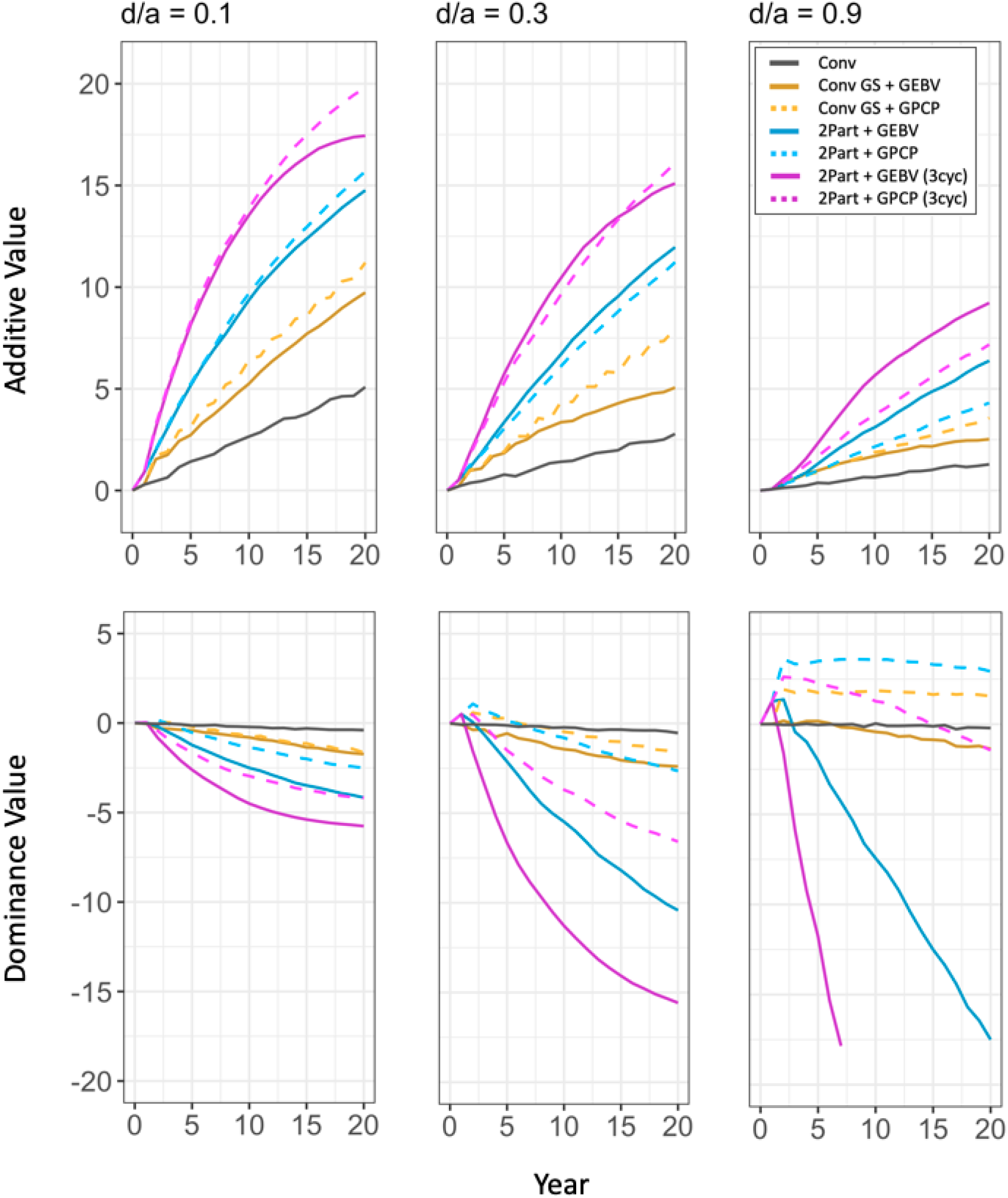
Additive values and the dominance values of the simulated breeding programs under different dominance degrees (d/a). In each of the upper panels (a-c), the additive values are plotted in clonal stage 1 for the future breeding phase. The lower panels (d-f) plot the dominance values. Each line shows the mean value for the 10 simulated replications. The different types of breeding program are shown in different colours. The conventional breeding program (Conv) is gray. The conventional breeding program with genomic selection (Conv GS) is yellow. The two-part breeding program with genomic selection (2Part) is shown in blue with one crossing cycle per year and in purple with three crossing cycles per year. The two types of parent selection were shown in different line-styles. Selection based on Genomic Estimated Breeding Value (GEBV) is shown by continuous lines. Selection based on Genomic Prediction of Cross Performance (GPCP) is shown by dashed lines. Additive values and dominance values at the beginning of the future breeding phase (year 0) were centred at zero.

**Figure 6.**
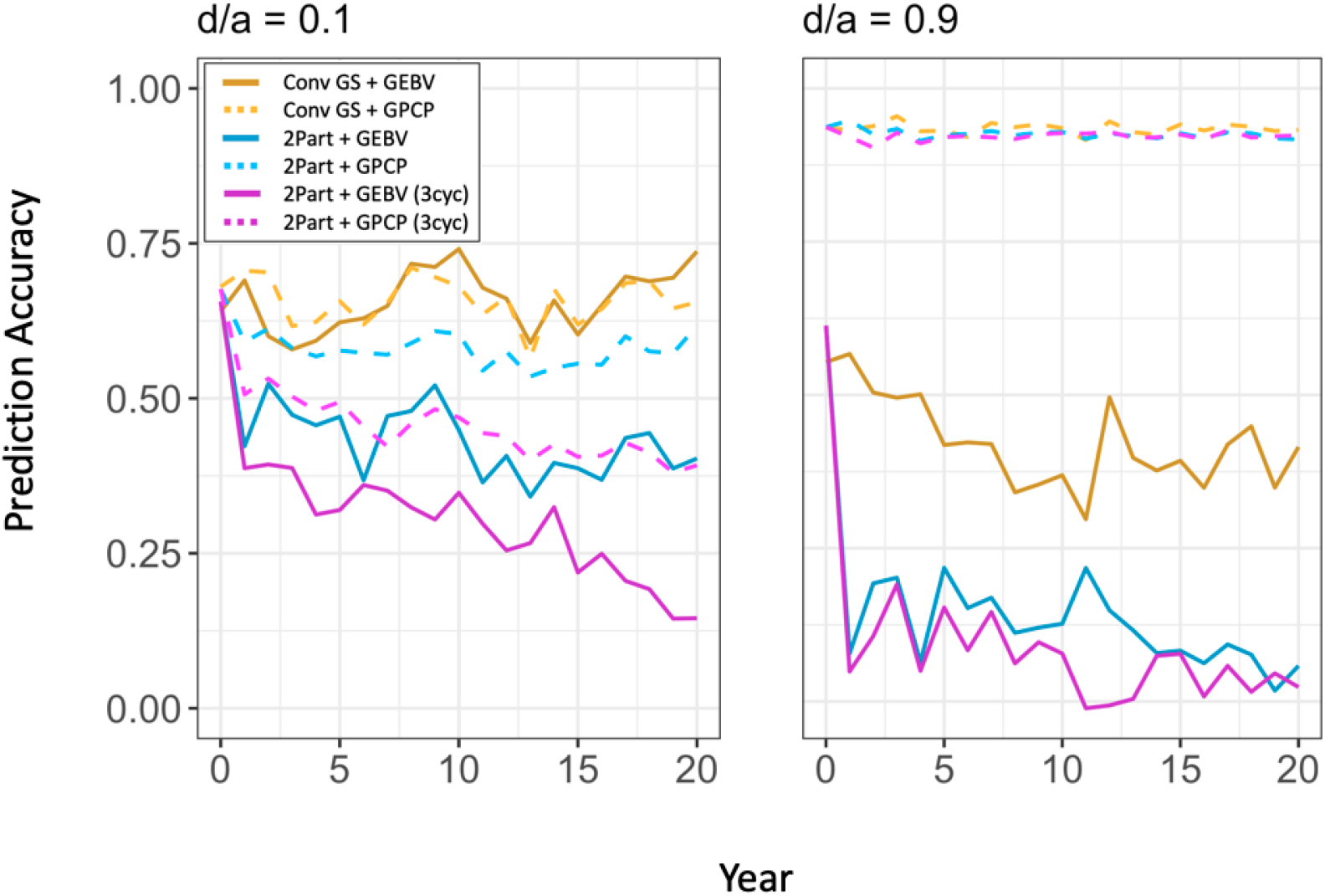
Prediction accuracy for selection of new parents under different dominance degrees (d/a). In each panel, prediction accuracy is plotted for the future breeding phase of the breeding programs with genomic selection. Each line shows the mean prediction accuracy for the 10 simulated replications. The different types of breeding program are shown in different colours. The conventional breeding program with genomic selection (Conv GS) is yellow. The two-part breeding program with genomic selection (2Part) is shown in blue with one crossing cycle per year and in purple with three crossing cycles per year. The two types of parent selection were shown in different line-styles. Selection based on Genomic Estimated Breeding Value (GEBV) is shown by continuous lines. Selection based on Genomic Prediction of Cross Performance (GPCP) is shown by dashed lines. Prediction accuracy was measured in the seedling stage for the two-part breeding programs and in clonal stage 1 for the conventional breeding program with genomic selection.

### Genetic gain

Selection of parents based on genomic predicted cross performance produced faster genetic gain than selection of parents based on genomic estimated breeding values unless dominance was negligible. This is shown in Figure 3, which plots genetic gain as the mean genetic value against time in clonal testing stage 1. The four panels show genetic gain under the different simulated dominance degrees for four types of breeding programs and two types of parent selection. As the dominance degree increased, selection of parents based on genomic prediction of cross performance produced increasingly more genetic gain than selection based on genomic estimated breeding values.

The three genomic selection breeding programs using genomic prediction of cross performance always produced more genetic gain than the conventional breeding program. The two variations of the two-part breeding program using genomic prediction of cross performance always produced the most genetic gain unless dominance was negligible (Fig. 3). However, while the two-part breeding program with three crossing cycles per year produced the most genetic gain when the dominance degree was 0.1 and 0.3, the two-part breeding program with one crossing cycle per year produced the most genetic gain when the dominance degree was 0.9. When the dominance degree was 0.1, the two-part breeding program gave 2.8 times the genetic gain of the conventional breeding program with one crossing cycle per year, and more than three times the genetic gain with three crossing cycles per year. When the dominance degree was 0.9, it gave almost 7 times the genetic gain of the conventional breeding program with one crossing cycle per year, and more than five times the genetic gain with three crossing cycles per year.

Figure 3 also shows that the two-part breeding program with parent selection based on genomic estimated breeding values and three crossing cycles per year generated the most genetic gain when the dominance degree was zero. However, after a sharp increase in the first few years, the rate of genetic gain drastically decreased and started to approach a plateau. The two-part breeding program with parent selection based on genomic estimated breeding values and one crossing cycle per year generated the second most genetic gain. In the first few years it showed a lower rate of genetic gain than both variations of the two-part breeding program using genomic prediction of cross performance. In the long term, however, both two-part breeding programs using genomic prediction of cross performance started to plateau and were outperformed by the two-part breeding program with parent selection based on genomic estimated breeding values and one crossing cycle per year.

Figure 3 also shows that selection of parents based on genomic estimated breeding values produced negative genetic gain over time when the dominance degree was high. All breeding programs showed a reduced rate of genetic gain when the dominance degree increased. However, this reduction was stronger when new parents were selected based on genomic estimated breeding values. Both variations of the two-part breeding program with parent selection based on genomic estimated breeding values produced even less genetic gain than the conventional breeding program when the dominance degree was 0.3 and 0.9. These results were not surprising as selection of parents based on genomic estimated breeding values gave a faster increase in the inbreeding coefficient than selection of parents based on genomic predicted cross performance when the dominance degree was high, which resulted in inbreeding depression.

### Genomic inbreeding coefficient

Selection of parents based on genomic predicted cross performance substantially reduced inbreeding when the dominance degree increased. This is shown in Figure 4, which plots the genomic inbreeding coefficient against time in clonal testing stage 1. The four panels show the inbreeding coefficient under the different simulated dominance degrees. As the dominance degree increased, all breeding programs showed a decreased growth rate of the genomic inbreeding coefficient. However, this decrease was much stronger when parents were selected based on genomic predicted cross performance compared to when genomic estimated breeding values were used.

Figure 4 also shows that the two-part breeding programs with selection of parents based on genomic predicted cross performance gave the strongest reduction in the genomic inbreeding coefficient as the dominance degree increased. When the dominance degree was zero, both breeding programs had almost approached complete inbreeding at the end of the future breeding phase. However, when the dominance degree was 0.9, the two-part breeding program with parent selection based on genomic predicted cross performance and one crossing cycle per year gave the lowest inbreeding coefficient, which was negative during the entire future breeding phase. The two-part breeding program with parent selection based on genomic predicted cross performance and three crossing cycles per year was also negative in the first half of the future breeding phase, but showed a slightly faster increase and became positive during the second half. By reducing the growth rate of the inbreeding coefficient when the dominance degree increased, selection of cross performance directly took the increasing importance of dominance effects to the total genetic value into account.

### Additive values and dominance values

Selection of parents based on genomic predicted cross performance enabled a better exploitation of the combined additive and dominance values than did selection of parents based on genomic estimated breeding values. This is shown in Figure 5, which plots the additive values and the dominance values against time in clonal testing stage 1. The three upper panels (a-c) show the additive values and the three lower panels (d-f) show the dominance values.

The two-part breeding program with parent selection based on genomic predicted cross performance and three crossing cycles per year gave the highest increase of the additive value over time when the dominance degree was 0.1 and 0.3 (Fig. 5a-c). The two-part breeding program with parent selection based on genomic estimated breeding values and three crossing cycles per year gave a lower additive value, as growth rate showed a stronger reduction over time and approached a plateau towards the end of the future breeding phase. However, when the dominance degree was 0.9, it gave the highest increase of the additive value.

Figure 5 a-c also shows that the rate of increase of the additive value over time was reduced as the dominance degree increased. All breeding programs gave a lower additive value under high dominance degrees compared to when the dominance degree was low. The conventional breeding program always gave the lowest increase of the additive value.

Selection of parents using genomic prediction of cross performance generated increased dominance values as the dominance degree increased (Fig. 5d-f). It gave a reduction of the dominance value when the dominance degree was 0.1, but a strong initial increase when the dominance degree was 0.9. The increase of the dominance value compensated for the reduced rate of increase of the additive value as the dominance degree increased. The two-part breeding program with parent selection based on genomic predicted cross performance and one crossing cycle per year gave the strongest increase. When the dominance degree was high, the two-part breeding program with one crossing cycle per year and the conventional breeding program with genomic selection maintained a relatively stable, positive dominance value over the entire future breeding phase. The two-part breeding program with three crossing cycles per year, however, showed a continuous reduction of the dominance value over time. It also showed a faster reduction than the other two breeding programs when the dominance degree was 0.1 and 0.3.

Selection of parents based on genomic estimated breeding values did not effectively exploit the dominance value as the dominance degree increased. This is also shown in Figure 5 d-f. Both variations of the two-part breeding program with parent selection based on genomic estimated breeding values generated reduced dominance values as the dominance degree increased. This reduction in the dominance value over time became more extreme as the dominance degree increased, and was greater than the increase in the additive value over time when the dominance degree was high.

### Prediction accuracy of the parent selection method

The advantage of using genomic predicted cross performance to select parents over using genomic estimated breeding values was not only due to a better simultaneous exploitation of the additive value and the dominance value, but also resulted from a substantially higher prediction accuracy when the dominance degree was high. At higher dominance degrees, selection of parents based on genomic predicted cross performance became more accurate and selection of parents based on genomic estimated breeding values became less accurate. This is shown in Figure 6, which plots the prediction accuracy of the parent selection methods against time. The two panels show prediction accuracy under the dominance degrees of 0.1 and 0.9 for the three types of genomic selection breeding programs and two types of parent selection method. Prediction accuracy of the parent selection method was measured in the seedling stage for the two-part breeding programs and in clonal testing stage 1 for the conventional breeding program with genomic selection. Prediction accuracy of genomic predicted cross performance became more similar in the three genomic selection breeding programs when the dominance degree increased.

### Prediction accuracy of the genetic value in the seedling stage

Prediction accuracy of the genetic value of the seedlings increased when the dominance degree was increased. Figure 7 plots the prediction accuracy of the genetic value in the seedling stage over time. The two panels show prediction accuracy under the dominance degrees of 0.1 and 0.9. The two-part breeding program with parent selection based on genomic estimated breeding values and one crossing cycle per year always showed the highest prediction accuracy. Prediction accuracy was lower when parents were selected based on genomic predicted cross performance compared to genomic estimated breeding values. It also was lower when three crossing cycles per year were used compared to one crossing cycle. The difference in prediction accuracy due to the number of crossing cycles per year, however, became smaller as the dominance degree increased. The conventional breeding program with genomic selection using genomic predicted cross performance to select parents showed the lowest prediction accuracies under all dominance degrees.

**Figure 7.**
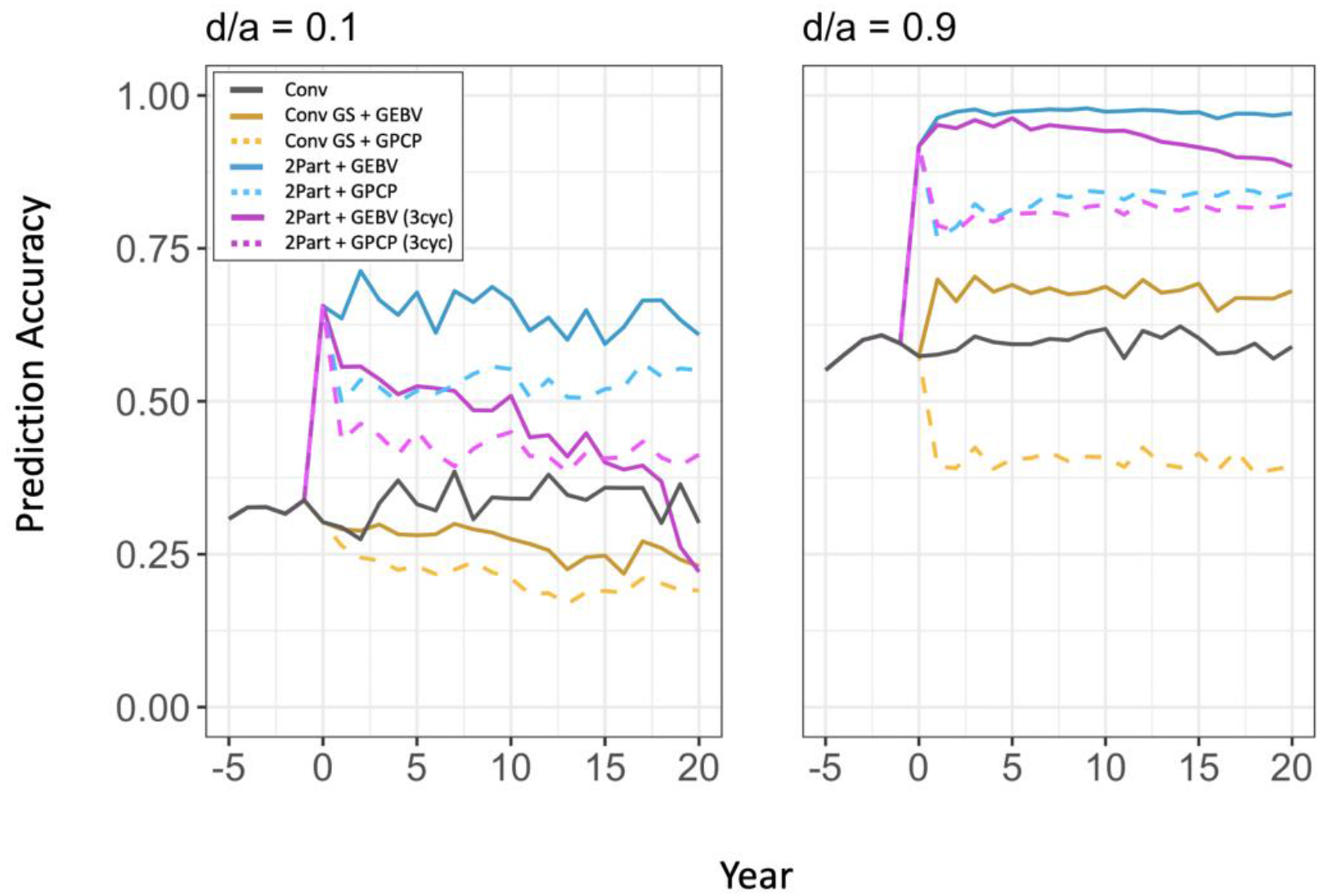
Prediction accuracy for the total genetic value of the seedlings under different dominance degrees (d/a). In each panel, prediction accuracy is plotted in the seedling stage for the entire burn-in breeding phase and the future breeding phase. Each line shows the mean prediction accuracy for the 10 simulated replications. The different types of breeding program are shown in different colours. The conventional breeding program (Conv) is gray. The conventional breeding program with genomic selection (Conv GS) is yellow. The two-part breeding program with genomic selection (2Part) is shown in blue with one crossing cycle per year and in purple with three crossing cycles per year. The two types of parent selection were shown in different line-styles. Selection based on Genomic Estimated Breeding Value (GEBV) is shown by continuous lines. Selection based on Genomic Prediction of Cross Performance (GPCP) is shown by dashed lines.

## Discussion

For genomic selection in clonal breeding programs to be effective, crossing parents should be selected based on genomic predicted cross performance unless dominance is negligible. To discuss this result, we first describe how genomic selection can improve clonal breeding programs under the assumption of additive genetic control. We show that the two-part breeding program enables effective exploitation of genomic selection in breeding clonally propagated crops. We also explain that under additive genetic control, differences in genetic gain between the two parent selection methods mainly resulted from an increased selection intensity when parents were selected based on genomic predicted cross performance compared to selection of parents based on genomic estimated breeding values. After the discussion of results when traits were under additive genetic control, we explain why genomic selection of new parents requires consideration of dominance effects unless dominance is negligible. We show that selection of parents based on genomic predicted cross performance enables efficient simultaneous exploitation of additive and dominance effects, which facilitates exploitation of pseudo-overdominance in the progeny of a cross when the dominance degree is high. We also show that multiple crossing cycles per year can have an adverse effect on long-term genetic gain, especially when the dominance degree is high. We then explain that, at higher dominance degrees, heterozygosity becomes a reliable predictor of the dominance value when parents are selected based on genomic predicted cross performance. Finally, we conclude that genomic prediction of cross performance could be an efficient method to select parents not only in clonal plant breeding programs, but also in other breeding programs for outbred individuals including animal breeding programs.

### Genomic selection of new parents improved genetic gain under additive genetic control

Under additive genetic control, genomic selection of new parents always produced faster genetic gain than phenotypic selection of new parents. This was observed regardless of whether parents were selected based on genomic estimated breeding values or based on genomic predicted cross performance.

As expected, the implementation of genomic selection improved the conversion of genetic variance into genetic gain in both variations of the two-part breeding program with one and three crossing cycles per year, respectively, and in the conventional breeding program with genomic selection. This improvement resulted from a shortened generation interval and an increased selection accuracy in early selection stages. As a consequence, the breeding programs with genomic selection also showed an accelerated depletion of genetic variance over time compared to the conventional breeding program (Fig. S1). This depletion was most severe when three crossing cycles per year were used, and it caused genetic gain to approach a plateau in the second half of the future breeding phase.

Our findings under additive genetic control were consistent with those of Gaynor et al. (2017) who used stochastic simulations to evaluate genomic selection strategies in plant breeding programs for developing inbred lines. We refer the reader to this study for a detailed description of the relationship between the generation interval, prediction accuracy and genetic variance when additive genetic control is assumed.

### A two-part breeding programs better exploit genomic selection than the conventional breeding program with genomic selection under additive genetic control

The two-part breeding programs enabled the best possible exploitation of genomic selection under additive genetic control. They produced between 2.3 times the genetic gain of the conventional breeding program when used with parent selection based genomic predicted cross performance and three crossing cycles per year, and three times the genetic gain of the conventional breeding program when used with parent selection based genomic estimated breeding values and three crossing cycles per year. The increased rates of genetic gain compared to the conventional breeding program resulted from a very short generation interval and an improved selection accuracy in the seedling stage.

Selection in the seedling stage poses a major challenge in clonal breeding programs due to a high selection intensity combined with low selection accuracy (Grüneberg et al., 2009; Bradshaw, 2016). The two-part breeding programs improved selection accuracy by replacing phenotypic selection with genomic selection, which enabled improvements in the selection criterion for seedlings. When phenotypic selection was used, seedlings were selected based on their observed *per se* performance. When genomic selection was used, seedlings were selected based on their predicted performance as clones because the genomic selection model was trained using data from the clonal testing stages.

Using genomic selection in the seedling stage improved selection accuracy for two reasons:

i. The phenotypic records in the clonal stages which were used to train the selection model had a higher heritability than the phenotypic records of the unreplicated seedlings.
ii. Marker alleles were replicated within and across years.

This increase of the selection accuracy also laid the foundation for the selection of new parents in the seedling stage, allowing for one or multiple cycles of crossing per year to minimize the length of the breeding cycle.

The conventional breeding program with genomic selection gave 1.7 times the genetic gain of the conventional breeding program when parents were selected based on genomic estimated breeding values and 1.9 times the genetic gain when parents were selected based on genomic predicted cross performance. Genomic selection was applied in clonal testing stage 1 and selection in the seedling stage was based on phenotypic *per se* performance. Hence, selection accuracy in the seedling stage was not improved compared to the conventional breeding program. The increased rate of genetic gain mainly resulted from a shortened generation interval and an improved selection accuracy in clonal testing stage 1.

### Selection of parents based on genomic predicted cross performance increased selection intensity compared to selection of parents based on genomic estimated breeding values under additive genetic control

Under additive genetic control, differences in genetic gain between the two parent selection methods mainly resulted from an increased selection intensity when parents were selected based on genomic predicted cross performance compared to selection of parents based on genomic estimated breeding values.

When genomic estimated breeding values were used, the 30 best genotypes were selected and randomly crossed to mimic a “good by good” crossing scheme. When genomic predicted cross performance was used, parents were selected based on the predicted mean genetic values of the F_1_ of a bi-parental cross. Under additive genetic control, the predicted mean genetic value of the F_1_ is equal to the mean genomic estimated breeding value of both parents. Selection of parents based on genomic predicted cross performance resulted in the excessive use of a few very good parents in many crosses. As a consequence, the selection intensity for parents was higher compared to when parents were selected based on genomic estimated breeding values and randomly crossed.

In the conventional breeding program with genomic selection, this increase in selection intensity resulted in more genetic gain over time compared to when parents were selected based on genomic estimated breeding values. In the two-part breeding programs, it resulted in more genetic gain in the first years, but thereafter genetic gain reached a plateau due to a depletion of genetic variance. This depletion of genetic variance was more severe when three crossing cycles per year were used.

A crossing strategy in a practical breeding program would probably lie somewhere in between the two simulated parent selection methods. A breeder would not randomly select crosses, but rather combine parents that are expected to generate improved progeny. Although very good genotypes may be used at high frequency, a breeder would make sure that an overly excessive use did not restrict the genetic variation in long-term.

### Genomic selection of new parents requires consideration of dominance effects unless dominance is negligible

If dominance is appreciable, genetic gain becomes a function of combined additive and non-additive gene action. If epistasis is ignored, the non-additive gene action is entirely determined by dominance. Achieving a high rate of genetic gain then depends on an efficiently balanced exploitation of the additive and dominance effects (Bradshaw, 2016).

In particular, this requires two opposed actions:

i. The frequency of alleles with beneficial additive genetic effects in homozygous state has to be increased to improve the additive value in the breeding population.
ii. Heterozygosity has to be maintained to exploit dominance effects and keep the dominance value at a high level in the breeding population.

While inbreeding can be deliberately used to increase the frequency of beneficial alleles in homozygous state and hence to improve the additive value, it also results in a reduction of heterozygosity and the dominance value. In the worst case scenario, the decrease in the dominance value over time will exceed the increase in the additive value, and the rate of genetic gain will become negative due to inbreeding depression. Hence, it is obvious that both the contribution of the additive value and the contribution of the dominance value to genetic gain must be taken into account when selecting the crossing parents of the next generation.

More specifically, this selection process requires a balanced exploitation of the additive value and the dominance value based on the dominance degree. As the dominance degree increases, the importance of the dominance value relative to the additive value also increases, indicating that heterozygosity should be conserved more effectively. Optimally, a method to select new parents would automatically balance the contribution of the additive and dominance components to sustain long-term genetic gain without any prior knowledge about the dominance degree.

### Selection of parents based on genomic predicted cross performance enabled an efficient simultaneous exploitation of additive effects and dominance effects

Selection of parents based on genomic prediction of cross performance enabled an efficient simultaneous exploitation of additive effects and dominance effects by reducing the increase in inbreeding over time when the dominance degree increased. This became a crucial feature to increase genetic gain over time when the dominance degree was high.

As the dominance degree increased, selection of parents based on genomic prediction of cross performance produced increasingly more genetic gain than selection based on genomic estimated breeding values. By definition, the genomic estimated breeding value is the sum of the average effects of the marker alleles of a genotype. These average effects are predicted for all markers simultaneously by performing a linear regression of the phenotypes in the training population on the marker genotypes, the concept described by Falconer (1985) for a one-locus model. Although the genomic estimated breeding value thereby generally captures a large part of the dominance interaction (Falconer and Mackay, 1996; Hill et al., 2008), this population-based predictor of the value of an individual parent for the progeny generation ignores dominance deviation.

In contrast, selection of parents based on genomic predicted cross performance fully captures both additive and dominance marker effects. It thereby enables prediction of the expected genetic value of the progeny of a certain cross rather than prediction of the value of an individual parent. The inclusion of non-additive effects can facilitate an enhancement and an improved exploitation of non-additive genetic variation compared to parent selection based on genomic estimated breeding values (Varona et al., 2018).

When parents were selected based on genomic predicted cross performance, the enhancement of non-additive genetic variation was a direct outcome of the reduced increase in inbreeding over time. The improved exploitation of non-additive genetic variation resulted from the efficiently balanced exploitation of the additive and dominance value when dominance was appreciable.

Interestingly, the genomic prediction model for cross prediction autonomously assigned more weight to the dominance value as dominance increased without any prior knowledge about the dominance degree. This was achieved by including a covariate associated with genomic inbreeding (heterozygosity) in the model, which accounted for directional dominance, and can be seen as an estimator for inbreeding depression caused by genomic inbreeding (Xiang et al., 2016; Varona et al., 2018). As the dominance degree increased, the value of crosses which maintained heterozygosity in the population increased, and genomic prediction of cross performance accurately predicted those crosses.

### Selection of parents based on genomic predicted cross performance enabled exploitation of pseudo-overdominance in the progeny of a cross when the dominance degree was high

The two-part breeding programs with parent selection based on genomic estimated breeding values gave negative genetic gain due to severe inbreeding depression when the dominance degree was high. After the first year of future breeding, the decrease in the dominance value over time was consistently higher than the increase in the additive value.

At first sight, this might seem surprising as we did not simulate overdominance. Under the one-locus model with a dominance degree < 1, the allelic combination with the beneficial allele in homozygous state will result in the highest genetic value of all pairwise allelic combinations. In this case, selection of parents based on the genomic estimated breeding value would be an efficient strategy to increase the frequency of the beneficial allele in the population over time, and hence to increase genetic gain. Only under overdominance does the heterozygote become superior to both homozygotes and therefore the fixation of the allele with the higher additive value would result in a reduction of the genetic value (Falconer and Mackay, 1996)

Overdominance seems to be an extremely rare phenomenon in nature. However, due to linkage disequilibrium (LD), haplotype blocks are the units of genetic transmission rather than single loci. When haplotype blocks with favourable alleles in repulsion phase are combined during sexual recombination, the cumulative effect of these loci can create pseudo-overdominance although the dominance degree at each locus is < 1 (Bingham et al., 1994; Bingham, 1998).

Selection of parents based on the genomic estimated breeding value will drive an increase in the frequency of the haplotype blocks with the highest sum of average effects. The heterotic effects due to pseudo-overdominance, however, are reduced, or get lost, from one generation to the next. Furthermore, even haplotype blocks with lower genomic estimated breeding values may contain beneficial alleles, which are removed from the population through selection. As a result, genetic variance is reduced, restricting long-term additive genetic gain.

Selection of parents based on genomic predicted cross performance, on the other hand, included the heterotic potential of a cross when predicting the performance of the progeny. In this way, non-additive effects due to complementation of haplotype blocks can be maintained in the population over several generations when their contribution to the genetic value is high. Furthermore, by preserving haplotype blocks with lower genomic estimated breeding values for a few generations, recombination will make the beneficial alleles that they contain available for sustainable long-term genetic gain.

### Multiple crossing cycles per year using genomic prediction of cross performance without updating the prediction model can have an adverse effect on long-term genetic gain especially when the dominance degree is high

In the two-part breeding programs with parent selection based on genomic predicted cross performance, genomic inbreeding increased faster with three crossing cycles per year compared to one crossing cycle per year. While using three crossing cycles per year gave more genetic gain than one crossing cycle when the dominance degree was low, it gave less genetic gain when the dominance degree was high.

As the dominance degree increased, maintaining a low level of inbreeding became crucial to ensure a sustainable, long-term exploitation of dominance effects. We hypothesize that two factors caused the regulation of the inbreeding coefficient to be less efficient with three crossing cycles per year compared to one crossing cycle per year:

i. A reduced number of seedlings per crossing cycle.
ii. An irregular updating of the prediction model for selection of new parents.

The increased number of crossing cycles per year in combination with a reduced number of crosses and seedlings per cross resulted in an accelerated removal of haplotype block diversity from the breeding population. To equalize annual costs, the size of the seedling population was reduced from 12,000 to 4,000 seedlings per cross with three crossing cycles per year. Hence, the population became more susceptible to genetic drift and dominance effects due to complementation of haplotype blocks could not be maintained over multiple generations.

The irregular updating of the prediction model for the selection of new parents resulted in a less efficiently balanced exploitation of additive and dominance effects. Although multiple cycles of crossing and selection per year effectively reduced the generation interval, the genomic prediction model was updated only once a year, and selection of new crosses became increasingly less efficient. Assuming purely additive gene action in a simulation of a line breeding program, Gaynor et al. (2017) found that the increased genetic distance between the training and prediction population caused selection accuracy to drop with every additional crossing cycle. Although we also observed a reduction in prediction accuracy with an increased number of cycles (Fig. S4), the unchanged weights assigned to additive and dominance effects by the prediction model contributed more strongly to the accelerated reduction of heterozygosity. While inbreeding increased with every crossing cycle, the covariate associated with genomic inbreeding in the prediction model remained unchanged for two more cycles and could not sufficiently counteract inbreeding. When the model was then updated again in the following year, the loss of heterozygosity could not be completely reversed, which became especially problematic at a high dominance degree.

These results indicate that genomic prediction of cross performance to maximize genetic gain in the progeny generation might not be the best method to select new parents when multiple cycles of crossing and selection per year are used. To solve this problem, we hypothesize that a strategy such as optimal contribution selection could be useful to maximize long-term genetic gain as shown by Gorjanc et al. (2017) in a two-part line breeding program with multiple crossing cycles per year.

### Heterozygosity became a reliable predictor of the dominance value when the dominance degree was high

Prediction accuracy of genomic predicted cross performance increased as the dominance degree increased. Furthermore, prediction accuracy of the genetic value of the seedling genotypes increased as the dominance degree increased. Both prediction criteria included a non-additive term in the prediction model to capture dominance effects.

We infer that marker-based heterozygosity became an accurate predictor of non-additive genetic effects for selection of new parents especially when the dominance degree was high. This was mostly driven by including the covariate associated with genomic inbreeding (heterozygosity) in the model, which accounted for directional dominance. The two-part breeding programs especially benefited from the increase in prediction accuracy when the dominance degree increased.

Not only could cross performance be predicted more accurately, but selection accuracy in the seedlings also was significantly increased under high dominance degrees. Both factors contributed to the two-part breeding programs with genomic predicted cross performance generating the most genetic gain over time when dominance was appreciable.

### Implications for other breeding programs for outbred individuals

We expect genomic predicted cross performance could be used in other breeding programs for outbred individuals, such as animal breeding programs, to increase rates of genetic gain. As with clonal crops, animal breeding programs must also account for the detrimental effects of inbreeding depression. Animal breeders use various strategies to accomplish this ranging from rule-of-thumb recommendations to avoid matings between close relatives to optimal contribution selection, a numeric technique for limiting population level inbreeding (Woolliams et al., 2015). We expect genomic predicted cross performance to outperform these techniques by directly estimating progeny performance and thereby accounting for inbreeding depression in a purely data-driven manner, given the prediction model is constantly updated. However, when multiple cycles of crossing and selection per year are used without updating the prediction model, genomic prediction of cross performance to maximize genetic gain in the progeny generation might not be the best method to select new parents. In this case, implementing a strategy like optimal contribution selection might be useful to maximize long-term genetic gain, outlining an important topic for further research.

## Acknowledgments

The authors acknowledge the financial support from Innovate UK (132748).

## References

Bassil, N.V., T.M. Davis, H. Zhang, S. Ficklin, M. Mittmann, et al. 2015. Development and preliminary evaluation of a 90 K Axiom® SNP array for the allo-octoploid cultivated strawberry Fragaria × ananassa. BMC Genomics 16(1). doi: 10.1186/s12864-015-1310-1.

Bingham, E.T. 1998. Role of Chromosome Blocks in Heterosis and Estimates of Dominance and Overdominance. In: Larnkey, K.R. and Staub, J.E., editors, CSSA Special Publications. Crop Science Society of America, Madison, WI, USA. p. 71–87

Bingham, E.T., R.W. Groose, D.R. Woodfield, and K.K. Kidwell. 1994. Complementary Gene Interactions in Alfalfa are Greater in Autotetraploids than Diploids. Crop Sci. 34(4): 823–829. doi: 10.2135/cropsci1994.0011183X003400040001x.

Bisognin, D.A. 2011. Breeding vegetatively propagated horticultural crops. Crop Breed. Appl. Biotechnol. 11(spe): 35–43. doi: 10.1590/S1984-70332011000500006.

Bradshaw, J. 2016. Plant breeding: past, present and future. Springer, Cham.

Chen, G.K., P. Marjoram, and J.D. Wall. 2009. Fast and flexible simulation of DNA sequence data. Genome Res. 19(1): 136–142. doi: 10.1101/gr.083634.108.

Crossa, J., P. Pérez-Rodríguez, J. Cuevas, O. Montesinos-López, D. Jarquín, et al. 2017. Genomic Selection in Plant Breeding: Methods, Models, and Perspectives. Trends Plant Sci. 22(11): 961–975. doi: 10.1016/j.tplants.2017.08.011.

van Dijk, T., G. Pagliarani, A. Pikunova, Y. Noordijk, H. Yilmaz-Temel, et al. 2014. Genomic rearrangements and signatures of breeding in the allo-octoploid strawberry as revealed through an allele dose based SSR linkage map. BMC Plant Biol. 14(1): 55. doi: 10.1186/1471-2229-14-55.

Falconer, D.S. 1985. A note on Fisher’s ‘average effect’ and ‘average excess.’ Genet. Res. 46(3): 337–347. doi: 10.1017/S0016672300022825.

Falconer, D.S., and T.F.C. Mackay. 1996. Introduction to quantitative genetics. 4. ed., [16. print.]. Pearson, Prentice Hall, Harlow.

Gaynor, R.C., G. Gorjanc, A.R. Bentley, E.S. Ober, P. Howell, et al. 2017. A Two-Part Strategy for Using Genomic Selection to Develop Inbred Lines. Crop Sci. 57(5): 2372–2386. doi: 10.2135/cropsci2016.09.0742.

Gaynor, R.C., G. Gorjanc, D. Wilson, and J.M. Hickey. 2019. AlphaSimR: Breeding Program Simulations.

Gemenet, D.C., and A. Khan. 2017. Opportunities and Challenges to Implementing Genomic Selection in Clonally Propagated Crops. In: Varshney, R.K., Roorkiwal, M., and Sorrells, M.E., editors, Genomic Selection for Crop Improvement. Springer International Publishing, Cham. p. 185–198

Goddard, M. 2009. Genomic selection: prediction of accuracy and maximisation of long term response. Genetica 136(2): 245–257. doi: 10.1007/s10709-008-9308-0.

Goddard, M.E., and B.J. Hayes. 2007. Genomic selection: Genomic selection. J. Anim. Breed. Genet. 124(6): 323–330. doi: 10.1111/j.1439-0388.2007.00702.x.

Gorjanc, G., R.C. Gaynor, and J.M. Hickey. 2017. Optimal cross selection for long-term genetic gain in two-part programs with rapid recurrent genomic selection. bioRxiv: 227215. doi: 10.1101/227215.

Grüneberg, W., R. Mwanga, M. Andrade, and J. Espinoza. 2009. Selection methods. Part 5: Breeding clonally propagated crops. Plant Breed. Farmer Particip.: 275–322.

Hill, W.G., M.E. Goddard, and P.M. Visscher. 2008. Data and Theory Point to Mainly Additive Genetic Variance for Complex Traits. PLOS Genet. 4(2): e1000008. doi: 10.1371/journal.pgen.1000008.

Meuwissen, T., B. Hayes, and M. Goddard. 2016. Genomic selection: A paradigm shift in animal breeding. Anim. Front. 6(1): 6–14. doi: 10.2527/af.2016-0002.

Sargent, D.J., F. Fernandéz-Fernandéz, J.J. Ruiz-Roja, B.G. Sutherland, A. Passey, et al. 2009. A genetic linkage map of the cultivated strawberry Fragaria × ananassa and its comparison to the diploid Fragaria reference map. Mol. Breed. 24(3): 293–303. doi: 10.1007/s11032-009-9292-9.

Sargent, D.J., Y. Yang, N. Šurbanovski, L. Bianco, M. Buti, et al. 2016. HaploSNP affinities and linkage map positions illuminate subgenome composition in the octoploid, cultivated strawberry (Fragaria×ananassa). Plant Sci. 242: 140–150. doi: 10.1016/j.plantsci.2015.07.004.

Su, G., O.F. Christensen, T. Ostersen, M. Henryon, and M.S. Lund. 2012. Estimating Additive and Non-Additive Genetic Variances and Predicting Genetic Merits Using Genome-Wide Dense Single Nucleotide Polymorphism Markers. PLOS ONE 7(9): e45293. doi: 10.1371/journal.pone.0045293.

Varona, L., A. Legarra, M.A. Toro, and Z.G. Vitezica. 2018. Non-additive Effects in Genomic Selection. Front. Genet. 9. doi: 10.3389/fgene.2018.00078.

Woolliams, J.A., P. Berg, B.S. Dagnachew, and T.H.E. Meuwissen. 2015. Genetic contributions and their optimization. J. Anim. Breed. Genet. 132(2): 89–99. doi: 10.1111/jbg.12148.

Xiang, T., O.F. Christensen, Z.G. Vitezica, and A. Legarra. 2016. Genomic evaluation by including dominance effects and inbreeding depression for purebred and crossbred performance with an application in pigs. Genet. Sel. Evol. 48(1). doi: 10.1186/s12711-016-0271-4.

