## Supplementary figures for "Genomic selection strategies for clonally propagated crops"

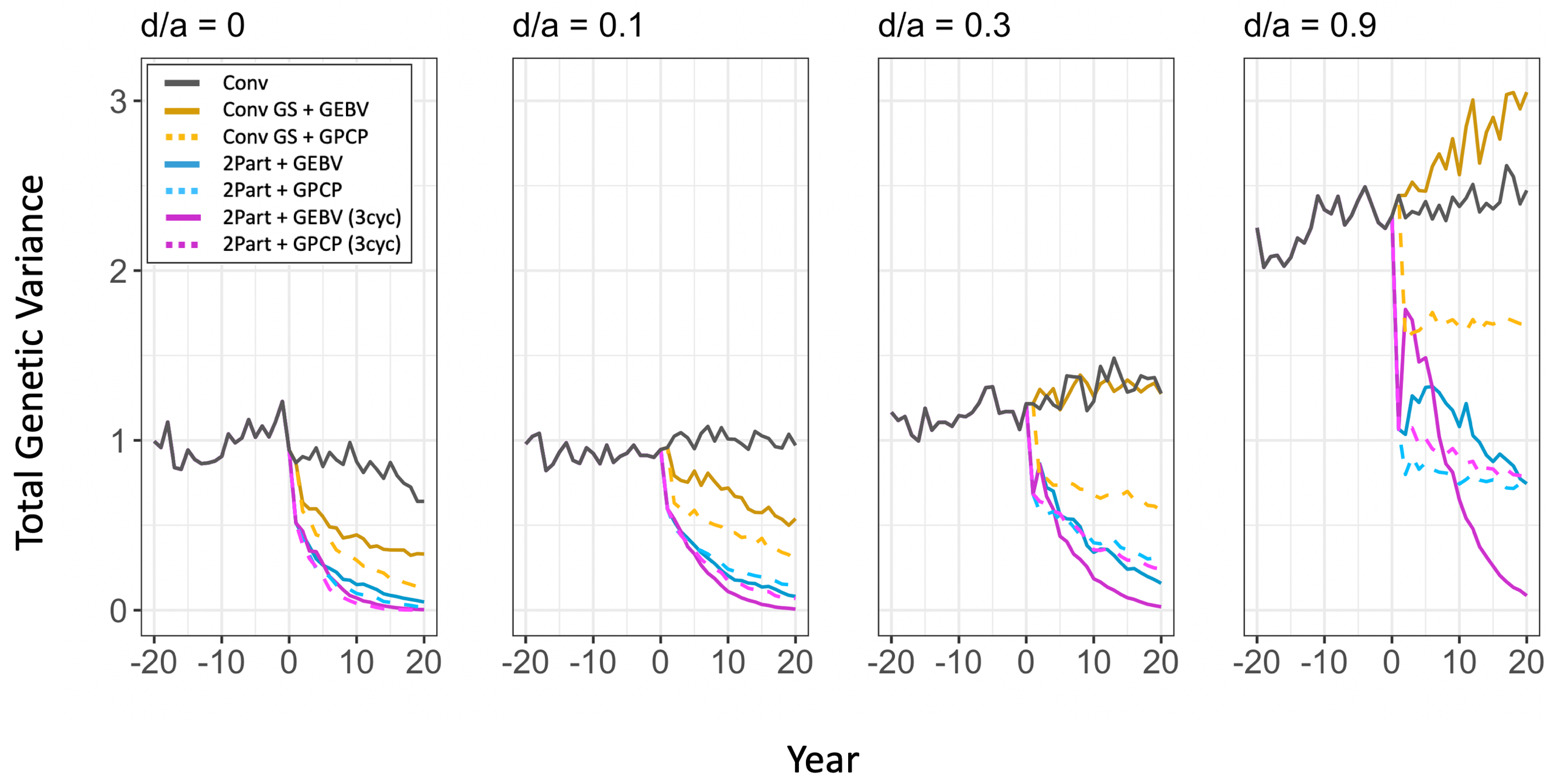

**Figure S1 Total genetic variance of the simulated breeding programs under different dominance degrees (d/a).** In each panel, total genetic variance is plotted in clonal stage 1 for the entire burn-in breeding phase and the future breeding phase. Each line shows the mean genetic variance for the 10 simulated replications. The different types of breeding program are shown in different colours. The conventional breeding program (Conv) is gray. The conventional breeding program with genomic selection (Conv GS) is yellow. The two-part breeding program with genomic selection (2Part) is shown in blue with one crossing cycle per year and in purple with three crossing cycles per year. The two types of parent selection were shown in different line-styles. Selection based on Genomic Estimated Breeding Value (GEBV) is shown by continuous lines. Selection based on Genomic Prediction of Cross Performance (GPCP) is shown by dashed lines.

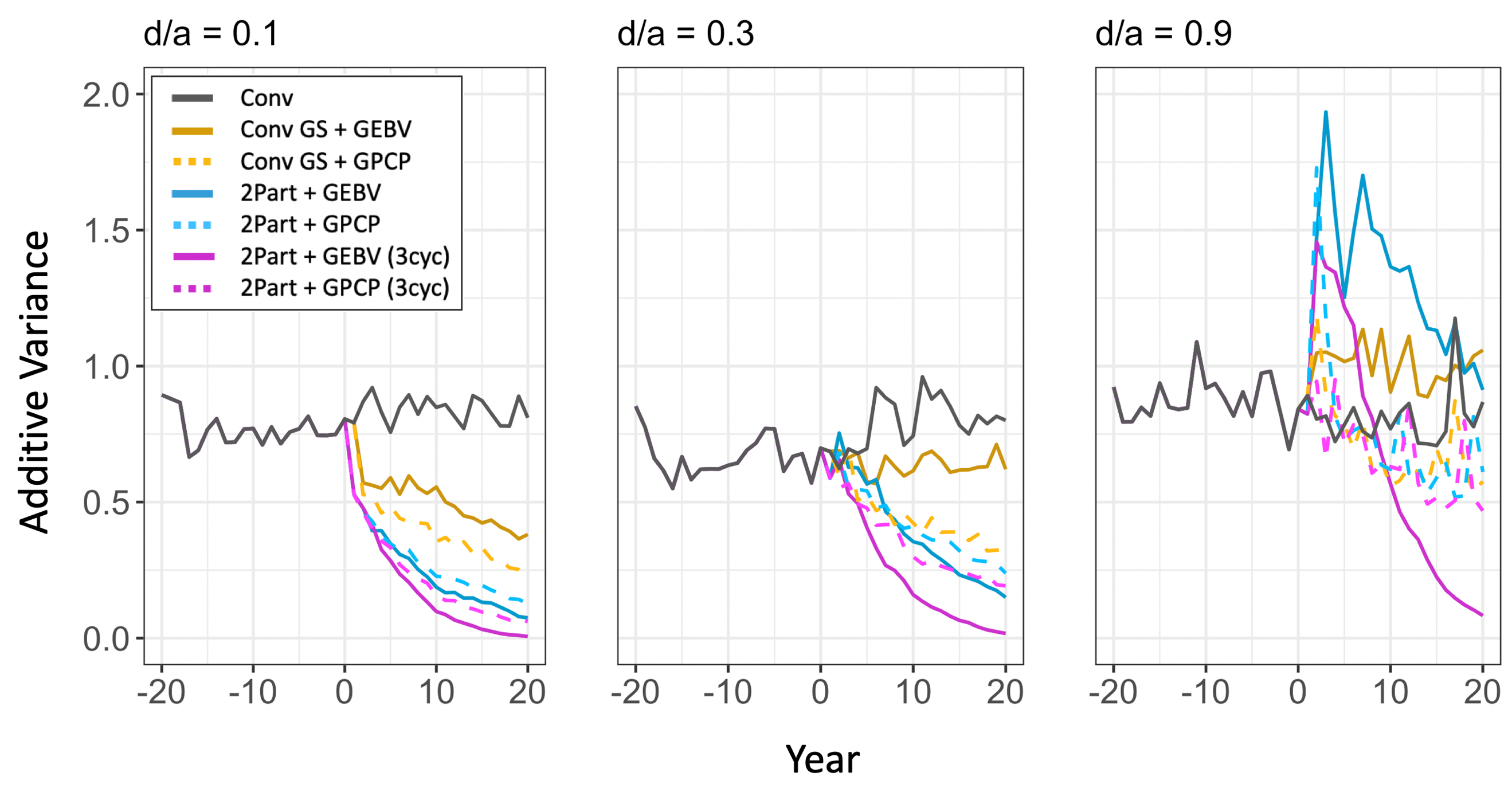

**Figure S2 Additive variance of the simulated breeding programs under different dominance degrees (d/a).** In each panel, additive variance is plotted in clonal stage 1 for the entire burn-in breeding phase and the future breeding phase. Each line shows the mean additive variance for the 10 simulated replications. The different types of breeding program are shown in different colours. The conventional breeding program (Conv) is gray. The conventional breeding program with genomic selection (Conv GS) is yellow. The two-part breeding program with genomic selection (2Part) is shown in blue with one crossing cycle per year and in purple with three crossing cycles per year. The two types of parent selection were shown in different line-styles. Selection based on Genomic Estimated Breeding Value (GEBV) is shown by continuous lines. Selection based on Genomic Prediction of Cross Performance (GPCP) is shown by dashed lines.

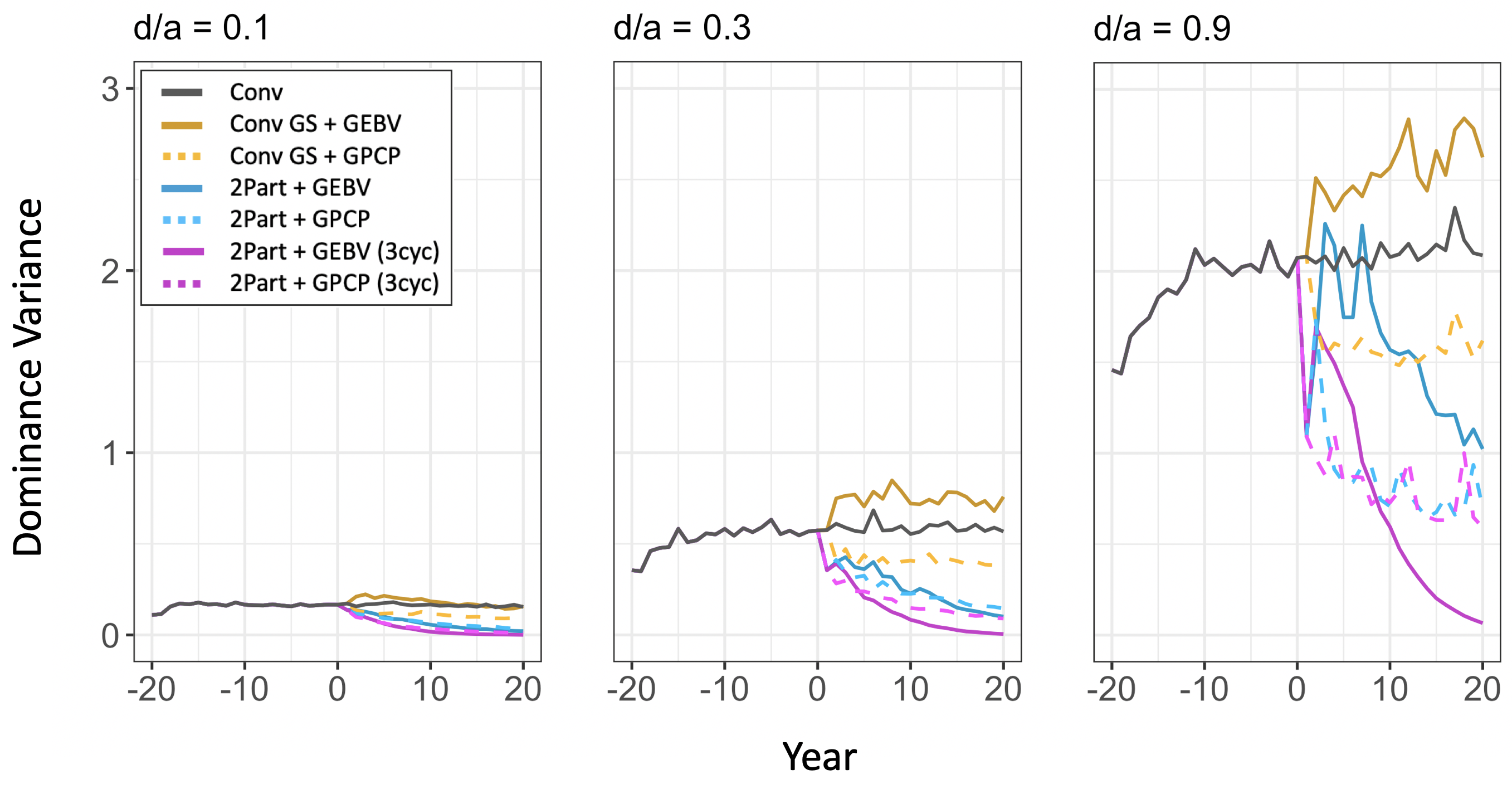

**Figure S3 Dominance variance of the simulated breeding programs under different dominance degrees (d/a).** In each panel, dominance variance is plotted in clonal stage 1 for the entire burn-in breeding phase and the future breeding phase. Each line shows the mean dominance variance for the 10 simulated replications. The different types of breeding program are shown in different colours. The conventional breeding program (Conv) is gray. The conventional breeding program with genomic selection (Conv GS) is yellow. The two-part breeding program with genomic selection (2Part) is shown in blue with one crossing cycle per year and in purple with three crossing cycles per year. The two types of parent selection were shown in different line-styles. Selection based on Genomic Estimated Breeding Value (GEBV) is shown by continuous lines. Selection based on Genomic Prediction of Cross Performance (GPCP) is shown by dashed lines.

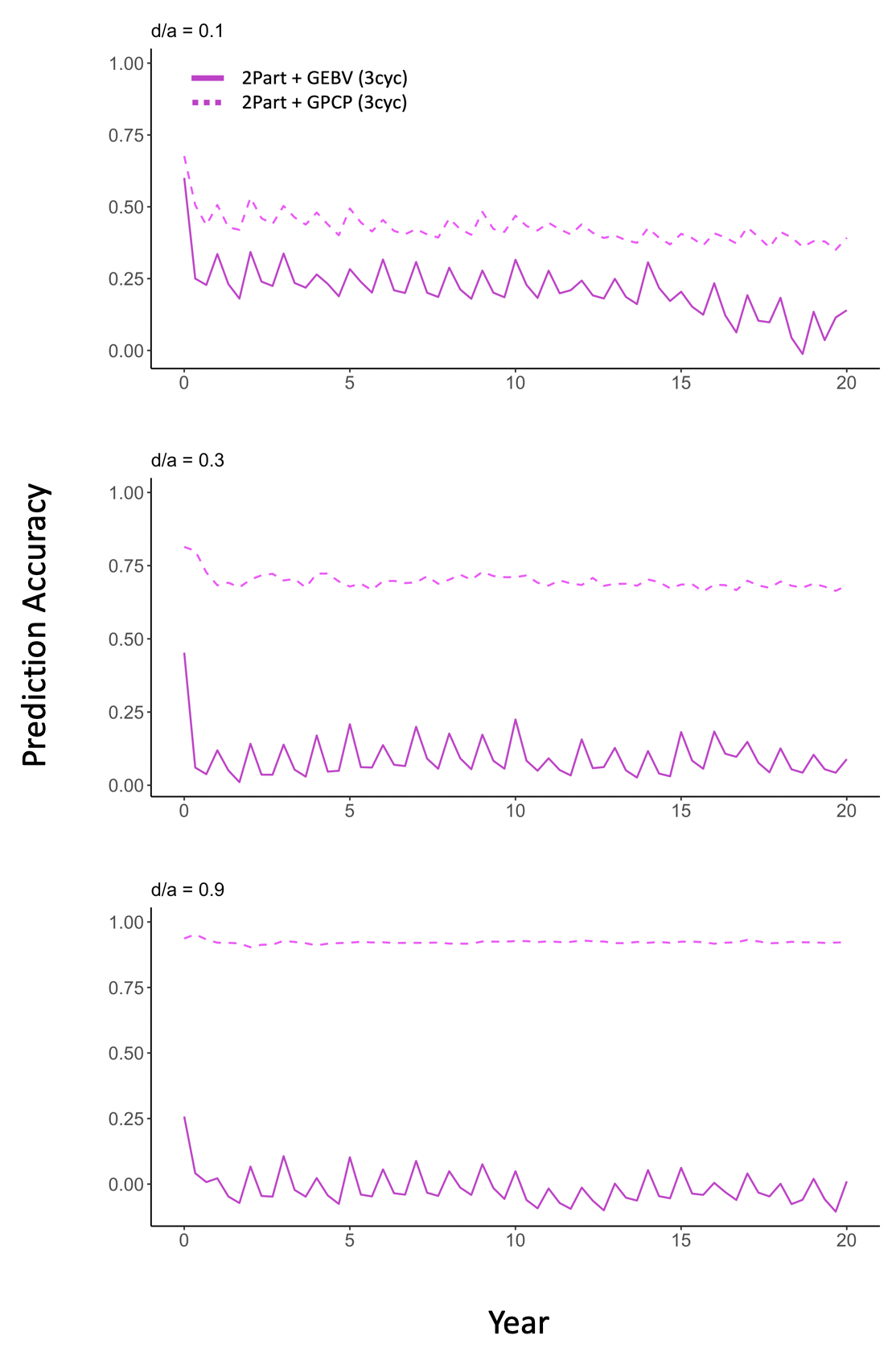

**Figure S4 Prediction accuracy for selection of new parents in the two-part breeding programs with three crossing cycles per year under different dominance degrees (d/a).** In each panel, prediction accuracy is plotted for the future breeding phase of the two-part breeding programs with three crossing cycles per year. Each line shows the mean prediction accuracy for the 10 simulated replications of the two breeding programs. The two types of parent selection were shown in different line-styles. Selection based on Genomic Estimated Breeding Value (GEBV) is shown by continuous lines. Selection based on Genomic Prediction of Cross Performance (GPCP) is shown by dashed lines. Prediction accuracy was measured in the seedling stage.
